# Early-life microbial intervention reduces colitis risk promoted by antibiotic-induced gut dysbiosis

**DOI:** 10.1101/2020.03.11.987412

**Authors:** Jun Miyoshi, Sawako Miyoshi, Tom O. Delmont, Candace Cham, Sonny T.M. Lee, Aki Sakatani, Karen Yang, Yue Shan, Evan Kiefl, Mahmoud Yousef, Sean Crosson, Mitchell Sogin, Dionysios A. Antonopoulos, A. Murat Eren, Vanessa Leone, Eugene B. Chang

## Abstract

Perturbations in the early life gut microbiome are associated with increased risk to complex immune disorder like inflammatory bowel diseases. We previously showed maternal antibiotic-induced gut dysbiosis vertically passed to offspring increases experimental colitis risk in IL-10 gene deficient (IL-10^−/−^) mice. While this could arise from emergence of pathobionts or loss/lack of essential microbes needed for appropriate immunological education, our findings suggest the latter. A dominant *Bacteroides* strain absent following antibiotic-induced perturbation was cultivated from murine fecal samples. Addition of this strain into mice with antibiotic-induced dysbiosis significantly promoted immune tolerance and reduced incidence of colitis in IL-10^−/−^ mice, but only if engrafted early in life, and not during adulthood. Thus, key members of the gut microbiome are essential for development of immune tolerance to commensal microbes in early life and their addition in presence of gut dysbiosis during this period can reduce colitis risk in genetically prone hosts.

**Highlights:**

- Specific gut microbes promote early life immune tolerance to key commensal microbes
- Loss of early life keystone microbes increases colitis risk in genetically prone hosts
- Emergence of absent commensal microbes late in life worsened colitis outcome
- Early life exposure to a missing keystone Bacteroides strain reduced colitis risk

## Introduction

Inflammatory bowel diseases (IBD) are chronic disorders of unclear etiology that arise from convergence of environmental, genetic, and microbial factors. As both prevalence and incidence of IBD have been increasing globally over the past century (Kaplan, 2015), changes in environmental factors including diet, hygiene, and lifestyle are hypothesized to alter gut microbiome-host factors to trigger IBD in genetically susceptible subjects (Bager et al., 2012; Benchimol et al., 2015; Frank et al., 2011; Gevers et al., 2014; Lewis et al., 2015; Machiels et al., 2014; Manichanh et al., 2006; Ott et al., 2004; Thia et al., 2008).

Recent population-based studies showed an association between antibiotic exposure in early life and increased risk for IBD (Hviid et al., 2011; Kronman et al., 2012; Ortqvist et al., 2019; Ungaro et al., 2014; Virta et al., 2012). These reports suggest perturbations of the gut microbiota at this life stage can increase risk for IBD by affecting the host immune system during a critical developmental window. However, human epidemiological studies have been unable to provide insights into causality and mechanism. Given that western diets, broad spectrum antibiotics, formula-feeding, and Caesarian section births have all been shown to impact the development of the early life gut microbiome (Broe et al., 2014; Hersh et al., 2011; Petersen et al., 2010; Stokholm et al., 2013), there is an unmet need to understand the potential consequences and the mechanistic underpinnings of these environmental changes on early life development processes such as immune and metabolic systems that are impacted by the gut microbiome. In this regard, we hypothesize that disease risk begins at a much earlier life stage when disturbances in developmental programming of immune and other systemic processes can have long lasting consequences and be difficult to reverse. Recognition of these problems during this developmental window is paramount as we work to develop effective interventions that ensure healthier outcomes and reduce disease risk.

To test this hypothesis, we used a well-established murine model of experimental colitis that recapitulates many epidemiological features of human IBD (Miyoshi et al., 2017). The model includes administration of a broad spectrum antibiotic cefoperazone (CPZ). This treatment creates gut microbiota disturbances during the peripartum period of interleukin (IL)-10 deficient dams which is then vertically transmitted to the offspring. The acquired gut microbiota perturbation then skews immune development, increasing spontaneous colitis and chemical-induced colitis risk and severity. In addition, a clear causal link between CPZ-induced gut microbiota perturbations and pro-inflammatory immune development are evident, findings that have recapitulated by another group using a similar approach (Schulfer et al., 2018).

Here, our goal was to gain mechanistic insights into how antibiotic perturbed microbiota drives aberrant early life immunological development resulting in increased IBD risk using our peripartum-CPZ model. The study was designed to answer the following questions: (1) Is there a window of immune development early in life that is critically dependent on proper gut microbiome development? (2) Does this window close later in life, resulting in long-term colitis risk, especially in genetically prone hosts? (3) If immune tolerance to missing commensal microbes does not develop, is late life exposure associated with increased colitis risk in genetically prone hosts? (4) Can proper immune development be restored with a single indigenous microbial strain absent in CPZ-induced gut perturbations to reduce risk of spontaneous colitis? and (5) When does this intervention have to take place?

Our findings provide proof-of-concept that perturbations in the early life gut microbiome can negatively impact immune development, increasing risk for disease in genetically susceptible hosts. To further investigate whether the long lasting effects of early life perturbation can be reversed through microbial therapies, we cultured a dominant *Bacteroides* population we have identified in our mice cohort using genome-resolved metagenomics. Our analyses showed that this strain (‘*Bacteroides* sp. CL1-UC’) is capable of engrafting into the dysbiotic gut microbiota after antibiotic treatment, altering subsequent microbial membership and significantly reducing risk for spontaneous colitis development, but only if the intervention is performed early in life. Without development of immune tolerance to missing gut microbes, later life exposure is associated with increased colitis risk in a genetically prone host. Together, our findings support the notion that perturbations in early life immune and gut microbiome development can significantly affect subsequent states of health and disease.

## Results

### Maternal peripartum antibiotic exposure and vertical transmission of the resulting perturbed microbiota promotes chronic imbalance of CD4^+^ T cell subsets in offspring of IL-10 deficient dams

Using the peripartum-CPZ exposure protocol in IL-10 deficient (IL-10 KO) mice, we previously reported the vertical transmission of CPZ-induced microbiota from dams to their offspring leads to skewing of CD4^+^ T cell subpopulations in pups at weaning (3 weeks of age) that was consistent with a shift towards Th1 (type 1 helper T cell) and Th17 (T helper 17 cell) development(Miyoshi et al., 2017). We employed an identical peripartum-CPZ treatment protocol as previously performed and harvested cells from spleen (SPLN) and mesenteric lymph nodes (MLNs) of IL-10 KO offspring at 7 weeks of age (Figure 1A). We assessed the number of 16S rRNA gene per gram of feces to estimate bacterial load and performed 16S rRNA gene amplicon sequencing analysis via Illumina MiSeq to assess α–diversity and β-diversity of the gut bacteriome. Male mice from CPZ-treated dams exhibited significantly reduced the estimated bacterial load at 3 weeks of age (weaning) compared to non-treated (NT) counterparts (*p* < 0.01) (Figure 1B). While the estimated bacterial load was restored by 7 weeks of age, Shannon diversity index remained lower in the CPZ group compared to NT mice (*p* < 0.01 at 3 and 7 weeks of age) (Figure 1C). Principal coordinate analysis (PCoA) plots for samples collected at 3 and 7 weeks of age are shown in Figure 1D. Unweighted UniFrac distances describe the operational taxonomical units (OTUs) existing in each sample, while weighted UniFrac distances take into account the proportional representation of the OTUs. Significant differences were observed in bacterial composition between the NT and CPZ groups at both 3 (unweighted *p* = 0.01 and weighted *p* < 0.05) and 7 weeks of age (unweighted *p* < 0.01 and weighted *p* < 0.01). Similar changes were observed in female animals (Figure S1A-C). Phylum level taxonomy are shown in Figure S1D. These findings are compatible with our previous report, confirming reproducibility of the CPZ treatment protocol. We examined live TCRβ^+^CD4^+^ cells (CD4^+^ T cells) expressing Foxp3 (regarded as regulatory T cells; Tregs), T-bet (Th1 cells), or RORγt (Th17 cells) at 7 weeks of age. Gating strategies for flow cytometry are shown in Figure S1E. Male CPZ group animals exhibited significant increases in Th1 cells and Th17 cells in SPLN (*p* < 0.05 for both), as well as Th17 cells in MLN (*p* < 0.05) compared to their NT counterparts (Figure 1E). Meanwhile, in female CPZ mice, MLN Tregs decreased significantly (*p* < 0.01) (Figure S1F). Overall, CPZ offspring presented with an aberrant balance of CD4^+^ T cells with lower percentages of Tregs with concomitant increases in percentages of Th1 and Th17 cells compared to NT offspring. This pro-inflammatory pattern at 7 weeks of age is similar to that observed at 3 weeks of age in our previous study (Miyoshi et al., 2017). Next, we harvested splenic DCs from NT and CPZ mice at 7 weeks of age followed by stimulation with feces obtained from age and sex-matched NT and CPZ animals. We observed no differences in DC IL-12p40 production between NT and CPZ mice following stimulation with vehicle, NT stool, or CPZ stool. Additionally, 7 weeks of age NT and CPZ feces elicited similar IL-12p40 production from NT and CPZ male and female DCs (Figures 1F and S1G, respectively). We posit the CPZ group immune imbalance is primarily manifested in adaptive rather than innate immunity. We also posit that the persistent imbalance of CD4^+^ T cell subsets observed in both male and female CPZ mice arises from lack of essential gut microbiome-derived cues required for proper immunological development during early life rather than from emergence of disease-promoting pathobionts. This notion is supported by our previous observation that conventionalization of GF IL-10 KO breeding pairs with CPZ-altered gut microbiota and subsequent vertical transfer to offspring is not associated with increased risk for spontaneous colitis (Miyoshi et al., 2017). As will be shown later, the findings of the present study add further support to this hypothesis.

**Figure 1.**
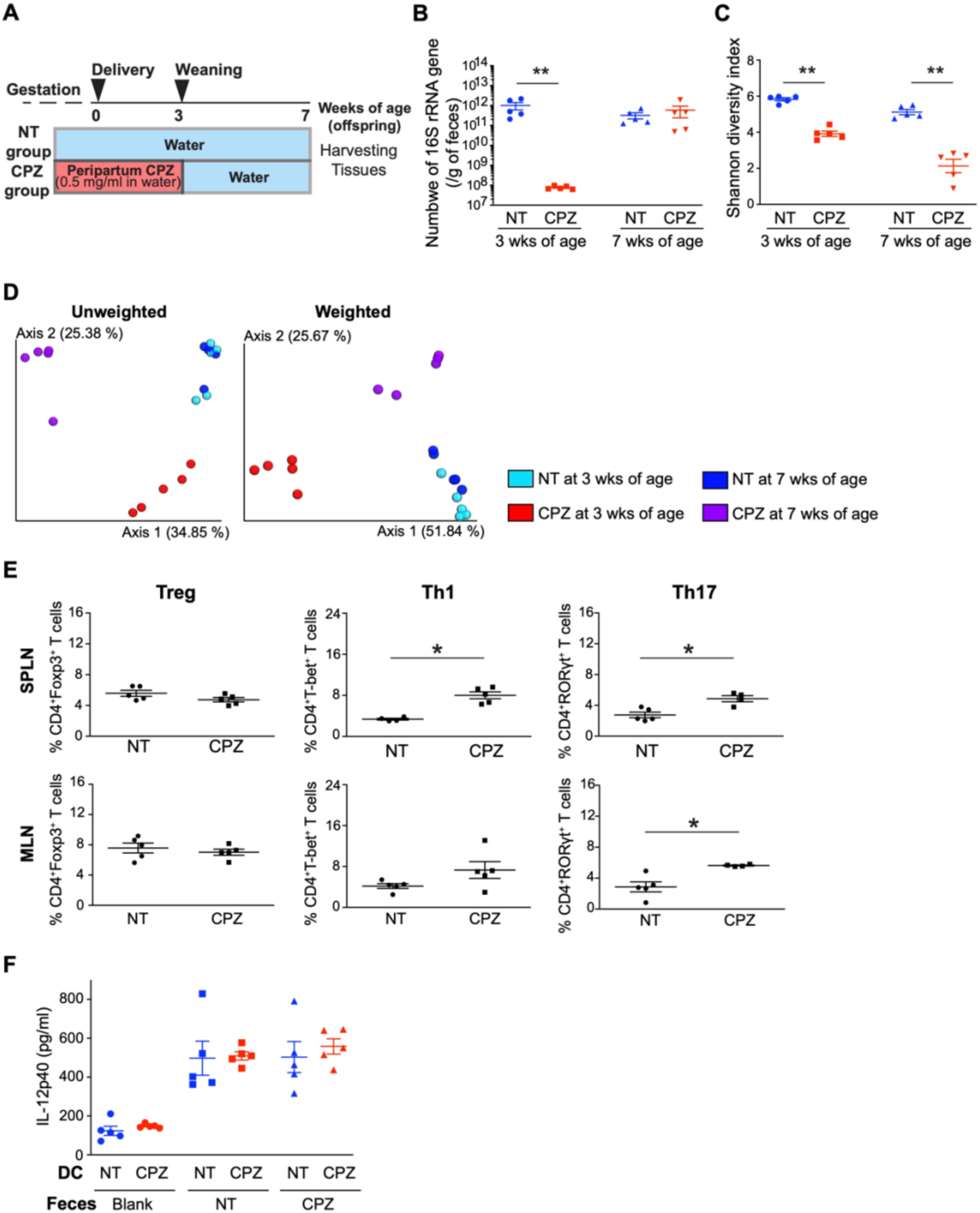
Maternal peripartum antibiotic exposure induces persistent gut microbiota perturbations and a lasting imbalance of CD4^+^ T cell subsets in offspring. (A) Study design using the IL-10 knock-out mouse model. CPZ group dams were treated with CPZ [(0.5 mg/ml) administration in drinking water] during pregnancy from day 14 of gestation until the end of the weaning period (3 weeks of age of offspring). (B) The number of 16S rRNA gene per gram of feces in male offspring from non-treated (NT) and CPZ-exposed dams at 3 and 7 weeks of age (*n* = 5 per group). (C) Shannon diversity index in male offspring at 3 and 7 weeks of age. (D) PCoA plots of both unweighted and weighted UniFrac distances of 16S rRNA gene amplicon sequences in NT and CPZ male pups at 3 and 7 weeks of age, respectively. (E) Flow cytometric analyses of live CD4^+^ T cells expressing Foxp3 (Treg), T-bet (Th1) or RORγt (Th17) in spleens (SPLNs) and mesenteric lymph nodes (MLNs) of NT versus CPZ males at 7 weeks of age (*n* = 4-5 per group). Data represent the percentage of live TCRβ^+^CD4^+^ cells. (F) IL-12p40 production of dendritic cells obtained from 7-week-old male NT (blue) and CPZ (red) pups with stimulation with fecal slurry (*n* = 5 per group). \**p* < 0.05 and \*\**p* < 0.01. Data represent mean ± SEM for (B), (C), (E), and (F). Female data are shown in the supplementary figure. See also Figure S1.

### CD4^+^ T cells from offspring of CPZ-treated IL-10 deficient dams promote a Th1 and Th17-inflammatory gut milieu

We next investigated if the skewed CD4^+^ T cell subpopulation balance that we observed exhibits a colitogenic impact *in vitro* and *in vivo*. We harvested both SPLN and MLN cells from CPZ and NT IL-10 KO offspring sacrificed at 7 weeks of age followed by non-specific stimulation with phorbol 12-myristate 13-acetate (PMA) and ionomycin. Male CPZ group animals exhibited significantly increased IFNγ producing CD4^+^ T cells in SPLN (*p* < 0.05) and significantly increased IL-17A producing CD4^+^ T cells in MLN (*p* < 0.05) compared to the NT group (Figure 2A). Female CPZ mice showed significantly more IFNγ and IL-17A producing CD4^+^ T cells in SPLN as compared to NT counterparts (*p* < 0.05 and *p* < 0.05, respectively) (Figure S2A). Gating strategies for flow cytometry are shown in Figure S2B. In addition to PMA and ionomycin, we also co-stimulated CD4^+^ T cells isolated from SPLNs and MLNs at 7 weeks of age with anti-CD3 and anti-CD28 antibodies. We again observed that MLN CD4^+^ T cells from male CPZ animals produced significantly more IL-17A compared to NT counterparts (*p* < 0.01) (Figure 2B). In females, significantly more IL-17A production was observed in SPLN CD4^+^ T cells from the CPZ group than the NT group (*p* < 0.01) (Figure S2C). Together, the results from these *in vitro* experiments suggest CPZ offspring exhibit a skewing in CD4^+^ T cells at 7 weeks of age that have a propensity to produce pro-inflammatory Th1 and Th17 cytokines, including IFNγ and IL-17A, as compared to those from NT offspring.

**Figure 2.**
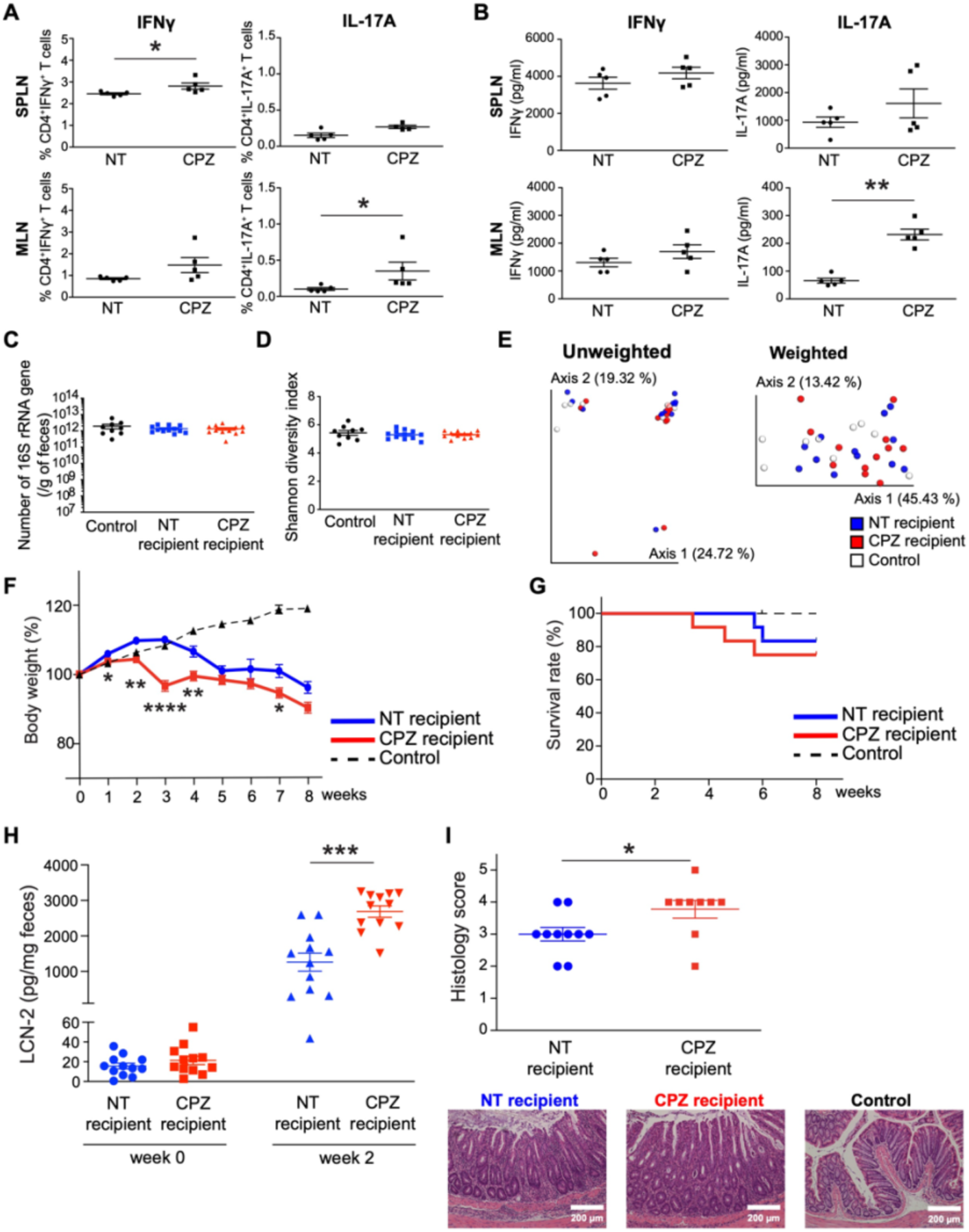
Maternal peripartum antibiotic exposure promotes colitogenic potential of CD4^+^ T cells in their offspring. (A) Flow cytometric analyses of live CD4^+^ T cells expressing IFNγ and IL-17A stimulated via PMA and ionomycin. Cells in spleens (SPLNs) and mesenteric lymph nodes (MLNs) were obtained from NT and CPZ males at 7 weeks of age. Data represent the percentage of live TCRβ^+^CD4^+^ cells (*n* = 5 per group). (B) IFNγ and IL-17A production of CD4^+^ T cells obtained from SPLN and MLN of 7-week-old NT and CPZ male pups with co-stimulation via anti-CD3 and anti-CD28 antibodies (*n* = 5 per group). (C) The number of 16S rRNA gene per gram of feces of male recipients before CD4^+^ T cell transfer (NT recipients: *n* = 12, CPZ recipients: *n* = 12, Control group: *n* = 9). (D) Shannon diversity index of male recipients before CD4^+^ T cell transfer. (E) PCoA plots of both unweighted and weighted UniFrac distances of 16S rRNA gene amplicon sequences in male recipients before CD4^+^ T cell transfer. (F) Percent weight change (expressed as % of starting weight) of male recipients after CD4^+^ T cell transfer. (G) Survival rate of male recipients during the 8-week observation period. (H) Fecal lipocalin-2 (LCN-2) levels before CD4^+^ T cell transfer (week 0) and at 2 weeks after post-transfer (week 2) in male recipients. (I) Histological assessment of colons from the surviving male recipients at week 8 (NT group: *n* = 10, and CPZ group: *n* = 9). Representative H&E histological sections of colon are presented for NT recipients, CPZ recipients, and the control group. \**p* < 0.05, \*\**p* < 0.01, and \*\*\**p* < 0.001. Data represent mean ± SEM for (A)-(D), (F), (H), and (I). Female data are shown in the supplementary figure. See also Figure S2.

### Adoptive transfer of CD4^+^ T cells from offspring of CPZ-treated IL-10 deficient dams promotes colitis development

To assess colitogenic potential of the skewed CD4^+^ T cell subsets *in vivo*, we transferred CD4^+^ T cells obtained from SPLNs of CPZ and NT IL-10 KO mice at 7 weeks of age into age- and sex-matched recombination activating gene 1 (RAG1) KO murine recipients. We prepared 2 groups; NT recipients with CD4^+^ T cells from NT IL-10 KO offspring and CPZ recipients with CD4^+^ T cells from CPZ IL-10 KO offspring. The control group received vehicle [phosphate buffered saline (PBS)] without CD4^+^ T cells. Since previous reports suggest that gut microbiota affects development and severity of colitis in this adoptive T cell transfer colitis model (Ostanin et al., 2009), we attempted to minimize gut microbiota variability among NT and CPZ RAG1 KO recipients prior to T cell transfer using our previously established and vetted protocol to minimize gut microbiota cage variance (Miyoshi et al., 2018). We confirmed that all groups of male (Figures 2C-E) and female (Figures S2D-F) RAG1 KO recipient mice harbored an identical number of 16S rRNA gene in feces (estimated bacterial load), Shannon diversity index, as well as exhibited no significant differences in unweighted and weighted UniFrac distances.

As expected, control PBS-injected mice did not exhibit body weight loss or other clinical symptoms of colitis. Male RAG1 KO CPZ recipients exhibited earlier and more severe body weight loss as compared to NT recipients (Figure 2F). The CPZ recipient survival rate was also decreased as compared to NT recipients (Figure 2G). Fecal lipocalin-2 (LCN-2), a non-invasive fecal colitis biomarker, exhibited more rapid elevation over time in CPZ recipients beginning 2 weeks after CD4^+^ T cell transfer (*p* < 0.001) (Figure 2H). The histological colitis score in animals that survived the entire 8-week post-CD4^+^ T cell transfer observation period was significantly higher in CPZ recipients as compared to NT recipients (Figure 2I). We observed similar findings in female recipients (Figure S2G-J). Overall, CD4^+^ T cells obtained from 7-week-old offspring of dams exposed to peripartum CPZ exhibited more colitogenic potential *in vivo* as compared to NT counterparts in both males and females. This finding supports the notion that early life immunological development is a determinant of health or colitis in genetically prone hosts.

### Identification and cultivation of a novel commensal Bacteroides strain that is essential for host immune development in early life

We explored microbes that may be playing a critical role in host immunological development in early life. Via 16S rRNA gene amplicon sequencing analysis, we previously demonstrated that many operational taxonomic units (OTUs) from the phylum *Bacteroidetes* were eradicated following maternal peripartum CPZ exposure and were also absent from CPZ-exposed offspring gut microbiota at certain time points post-weaning, including 3 and 7 weeks of age. However, some OTUs appeared to reemerge at 11 weeks of age, particularly in pups that developed spontaneous colitis later in life. It is notable that these reemerging OTUs appeared to be present as commensal bacteria within the NT maternal and offspring gut microbiota, yet none of the NT IL-10 KO dams or offspring developed colitis (Miyoshi et al., 2017). For a genome-resolved investigation of the microbial succession to identify microbial members that likely contribute to the colitis phenotype, we performed shotgun metagenomic sequencing followed by assembly and binning strategies to reconstruct metagenome-assembled genomes (MAGs). Metagenomic sequencing was performed on the same fecal DNA previously used for 16S rRNA amplicon sequencing (Miyoshi et al., 2017), including samples harvested from NT and peripartum CPZ exposed dams at weaning as well as 4 weeks after weaning, and NT and CPZ offspring at 3, 7, and 11 weeks of age (Supplementary Table S1 lists the details and accession IDs). No spontaneous colitis was observed in NT mice throughout the study while only some, but not all CPZ mice developed colitis at 12 to 23 weeks of age (mean age of colitis development was 16.5 weeks of age). However, neither NT nor CPZ offspring exhibited clinical symptoms of colitis at the time of fecal collections (3, 7, and 11 weeks of age). Along with NT samples, we included samples from CPZ pups that did (CPZ-colitis group) or did not (CPZ-no-colitis group) develop overt spontaneous colitis later in life. From the total set of 66 metagenomes, we reconstructed 216 non-redundant MAGs that were more than 70% and less than 10% redundant based on bacterial single-copy core genes and recruited on average 42.5% of reads from each metagenome (Table S1). Figure 3A displays the distribution of these MAGs across samples, where each column represents an individual MAG (216 in total) and each row represents an individual sample (66 in total). Analysis of MAGs via anvi’o (Eren et al., 2015) reveals gut bacterial composition at nearly the strain level.

**Figure 3.**
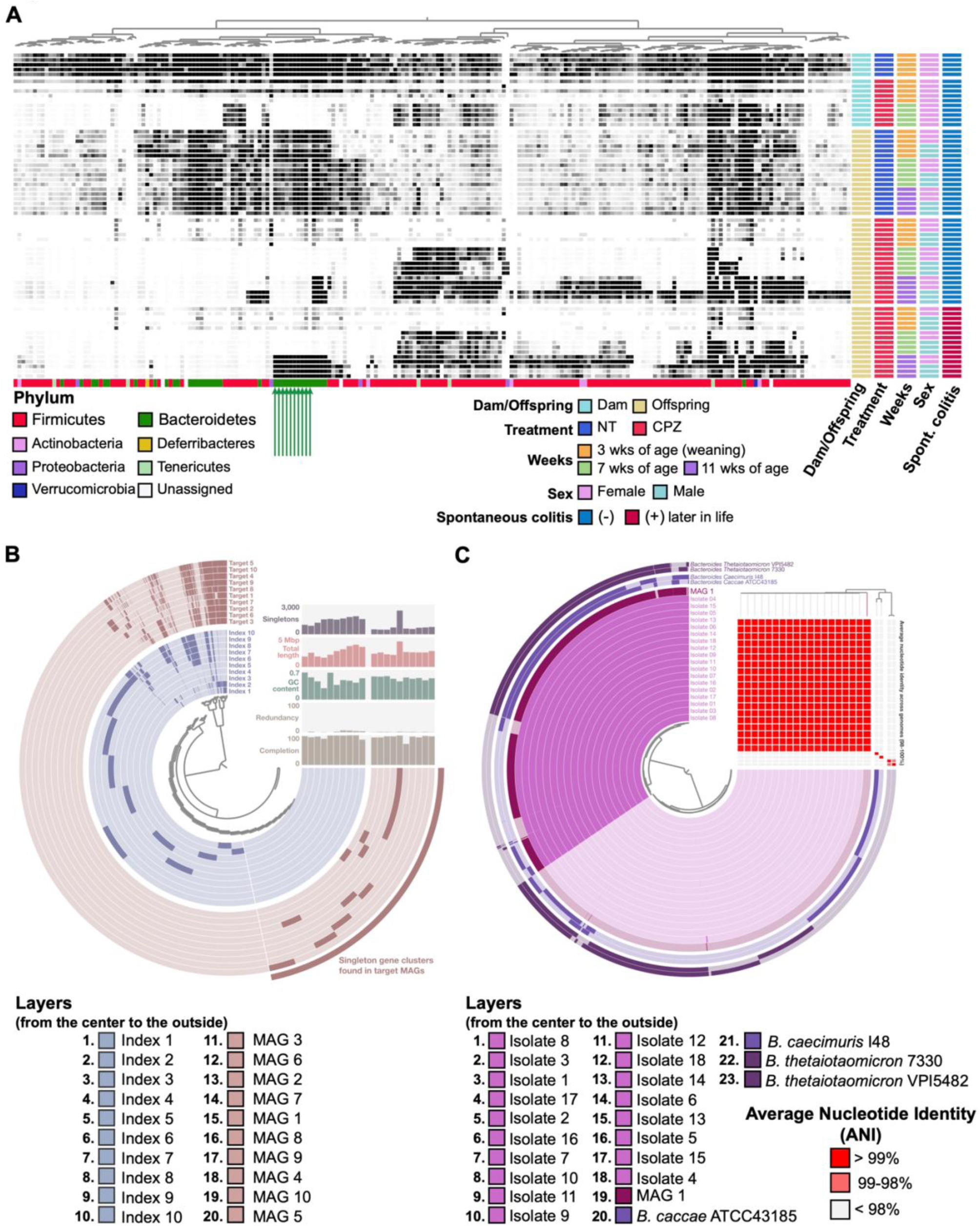
Identification and cultivation of a key commensal *Bacteroides* strain that is important for host immune development. (A) The detection rates of bacterial metagenomic-assembled genomes (MAGs) in fecal samples obtained from dams and pups over time. Each column represents a single MAG and each row represents an individual sample. The darker the cell, the greater the relative abundance. Metadata for each sample is presented to the right of the heat map [Dam/Offspring (origin of samples), Treatment (NT vs. CPZ), Weeks (pups at 3, 7 and 11 weeks of age), Sex (female vs. male), Spontaneous colitis (with/without development later in life)]. Colored bars at the bottom of the heatmap represent the MAG annotations at the phylum level. Green arrows indicate CPZ-sensitive MAGs that are residents of IL-10 KO dams and their NT offspring that did not develop colitis, but which emerged later in life of the CPZ group offspring that develop colitis. (B) Pangenome analysis to identify unique gene clusters in the target MAGs (MAGs 1-10), indicated by bottom green arrows in panel A. Each circular layer represents an individual MAG. The areas with deep color in each layer demonstrate the gene clusters contained by each MAG. The analysis of MAGs 1-10 with ten additional index MAGs is presented. Details of the MAGs are described in Table 1 and Table S2. (C) Pangenome analysis and assessment of average nucleotide identity (ANI) for 18 assembled genomes based on whole genome sequencing for 18 isolated cultivars compatible with MAG1 shown together with MAG1 and the four most similarly related complete genomes registered in the BLAST database. Each circular layer represents an individual MAG and darker areas in each layer show the gene clusters contained by each MAG contains. The ANI is presented in the right upper area corner in this figure. Each column represents an individual MAG corresponding to the circular layer. Each row presents an individual MAG in the order corresponding to the order of round layers from top to bottom. Red cell means > 99% identity. Detailed results of ANI are presented in Table S4. See also Figure S3.

We observed that 10 MAGs belonging to the phylum Bacteroidetes (MAG 1 to MAG 10; marked with green arrows in Figure 3A and details in Table 1) exhibited unique detection patterns: while they were clearly detected in NT dams and NT offspring at all time points, they were absent in dams and their offspring following peripartum CPZ exposure. Interestingly, these MAGs appeared to reemerge in the CPZ-colitis group but not in their CPZ-no-colitis group counterparts at 11 weeks of age in both males and females. These findings suggest that MAGs 1-10 represent bacterial populations that (1) are residents of our SPF IL-10KO mice, (2) exhibit sensitivity to CPZ, (3) appear to be non-pathogenic as they are maintained as commensal following the initial host exposure occurs early in life, and (4) can be colitogenic when initial exposure of the naïve, genetically susceptible host occurs late in life. These bacteria might play crucial roles during a critical period of early life for proper adaptive immune system development, e.g. CD4^+^ T cell subpopulations, and this absence could persist into adulthood with negative consequences in the host.

**Table 1.**
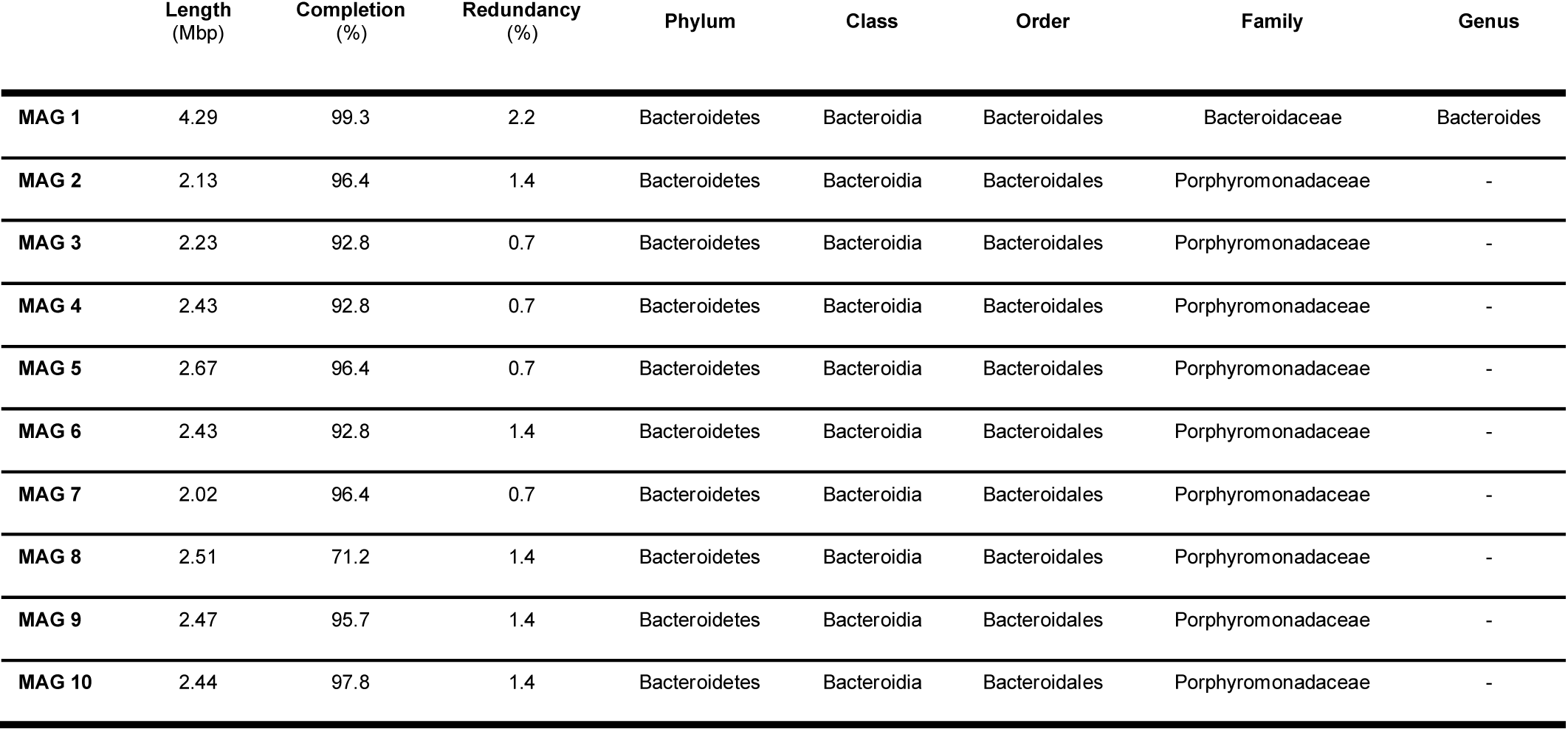
Metagenome-assembled genomes (MAGs) 1-10 information.

In order to further study the potential roles of these MAGs in both early life development of immune tolerance and later life immune system skewing and development of colitis, we sought to isolate resident bacterial strains corresponding to these MAGs from fecal samples obtained from 11 weeks old CPZ-colitis group male mice, a time point where their reemergence was observed. To ensure our cultivation efforts to obtain the target microbial members, we first used a pangenomics analysis strategy as implemented in anvi’o (Delmont and Eren, 2018) to design primers for each of MAGs of interest (MAGs 1-10) using gene clusters that are unique to them. In anvi’o pangenomics analysis, translated DNA sequences of predicted open reading frames are used to identify gene clusters in genome sequences. In addition to the 10 target MAGs, we also included in our pangenome an additional 10 index MAGs (Table S2) to minimize the likelihood of selecting a primer that is not unique to a single MAG (Figure 3B). Based on this analysis, we designed 4 targeted primer sets for MAGs 1-10, respectively (Table S3). Primers were validated using DNA from fecal samples containing MAG 1-10 genomes, i.e. fecal samples from 11-week-old CPZ-colitis group males used for metagenomic shotgun sequencing (Figure S3A). Targeted PCR was then performed on isolates cultivated anaerobically from the same fecal samples corresponding to MAGs 1-10. We subcultured an isolate that was positive for all 4 validated primer sets for each MAG and obtained 18 cultivars that appeared to be MAG 1. We were unable to establish cultivars for MAGs 2-10. This could, in part, be due to a significantly greater relative abundance of MAG 1 relative to MAGs 2-10, allowing MAG 1 to outcompete other similar strains under selective growth conditions. Whole genome sequencing (WGS) was performed on these 18 isolates to determine if they were indeed MAG 1, i.e. a single strain or if they were multiple strains with genomes that could not be distinguished with our 4 primer sets. Whole genomes for each isolate were assembled with PATRIC (Wattam et al., 2014) using raw sequence data. While the top 2 NCBI BLAST database hits for MAG 1 were *Bacteroides caccae* ATCC41385 and *Bacteroides caecimuris* I48, the top 2 BLAST hits for WGS assembled genomes for the 18 isolates were *Bacteroides thetaiotaomicron* 7330 and *Bacteroides thetaiotaomicron* VPI-5482. Both pangenomics and average nucleotide identity (ANI) analysis using anvi’o that included MAG 1, the 18 cultivar genomes, as well as the genomes of *B. caecimuris* I48, *B. thetaiotaomicron* 7330, *B. caccae* ATCC41385, and *B. thetaiotaomicron* VPI-5482 supported that the 18 WGS assembled genomes were indeed originated from a single strain (ANI 99.99 %) that was closely related to MAG 1 (ANI average 99.84%) and that these assembled genomes were distinct from the four *Bacteroides* species/strains based on BLAST results (ANI 81.32% to 90.17%) (Figure 3C and Table S4). Comprehensive genome analysis (CGA) via PATRIC (Davis et al., 2016; Edgar, 2004; Ondov et al., 2016; Stamatakis, 2014; Stamatakis et al., 2008; Wattam et al., 2014) revealed our isolate was most closely related to isolates from human feces, including *Bacteroides* sp. CAG:754 and reference organism *Bacteroides finegoldii* DSM 17565 (Figure S3B). However, our isolate was from murine feces and unique enough to be considered a novel strain, which we have provisionally named *Bacteroides* strain CL1-UC (Bc).

We next examined whether Bc exhibited immunogenic properties. Cultivars of *B. thetaiotaomicron* (Bt), *Lactobacillus murinus* (Lm), and *Lachnospiraceae bacterium* (Lb) were used as reference microbes. Bt was included because of its genomic similarity to Bc via BLAST. Lm and Lb were selected as representative CPZ-resistant microbes based on our metagenomic analysis. DCs from SPLN of 7 week old IL-10 KO mice were isolated and stimulated with lysates from these bacteria (Devkota et al., 2012). As shown in Figure S3C, IL-12p40 production was most significantly induced in DCs from both male and female mice following Bc lysate exposure. Bc lysate also stimulated more IL-12p40 and IFNγ production and promoted Th1 differentiation (Figure S3D) when DCs and naïve CD4^+^ T cells isolated from SPLN of 7 wk old IL-10 KO mice were cocultured. Thus, Bc, a member of the commensal bacterial consortium, exhibits unique immunogenic properties.

### Early life engraftment by Bacteroides CL1-UC (Bc) restores development of gut microbiota of offspring of CPZ treated dams

We investigated the impact of early life exposure to Bc on the gut microbiome of IL-10 KO offspring receiving CPZ-altered gut microbiota and on host immune development and colitis outcomes. The study was designed to answer the questions: (1) Can Bc engraftment into an early life CPZ-induced dysbiotic gut ecosystem restore immune development, even if it is eradicated later in life, reducing the risk of spontaneous colitis in IL-10 KO mice, i.e. immune imprinting?, and (2) Can Bc engraftment later in life (past the window of opportunity) restore commensal gut microbiota immune tolerance and reduce risk for development of spontaneous colitis in IL-10 KO mice? The first question was addressed by performing fecal microbiome transplantation (FMT) of CPZ-induced dysbiosis into male germ-free (GF) IL-10 KO mice at weaning (3 weeks of age). Donor fecal samples were collected from offspring of CPZ-treated dams where Bc was absent at 7-weeks-of-age. Immediately following FMT, for early engraftment, recipient mice received live Bc culture via gavage (100 µl of 5 × 10^8^ CFU/ml per day, two days) [designated early engraftment of Bc or EE group] or vehicle [no engraftment of Bc (NE) group] (Figures 4A and S4A). One week after gavage, PCR using Bc specific 16S rRNA gene markers showed that it successfully engrafted into 3 week old mice conventionalized with CPZ-associated microbiota, whereas it remained absent in non-engraftment (NE) controls who were not exposed to Bc (Figure S4B). The animals were tracked from 5 weeks of age (week 0) until 23 weeks of age (week 18) to assess health status, including body weight and clinical colitis scores (Figure 4A). At week 6 (11 weeks of age), all animals were treated with a cocktail of broad-spectrum antibiotic (Abx) protocol (including vancomycin, neomycin, and CPZ) applied in the drinking water for 72 hours for two purposes. First, to test if persistent engraftment by or just early exposure to Bc was sufficient to reduce risk for spontaneous colitis. Second, this arm of the study served as an additional control for studies presented in Figure 5. After a 24-hour recovery period, all mice were moved to new cages and gavaged with sterile PBS with 20% glycerol (Figures 4A and S4A). One week after Abx exposure, Bc had been eradicated in the EE group, despite recovery of the overall microbiota community (Figure S4C).

**Figure 4.**
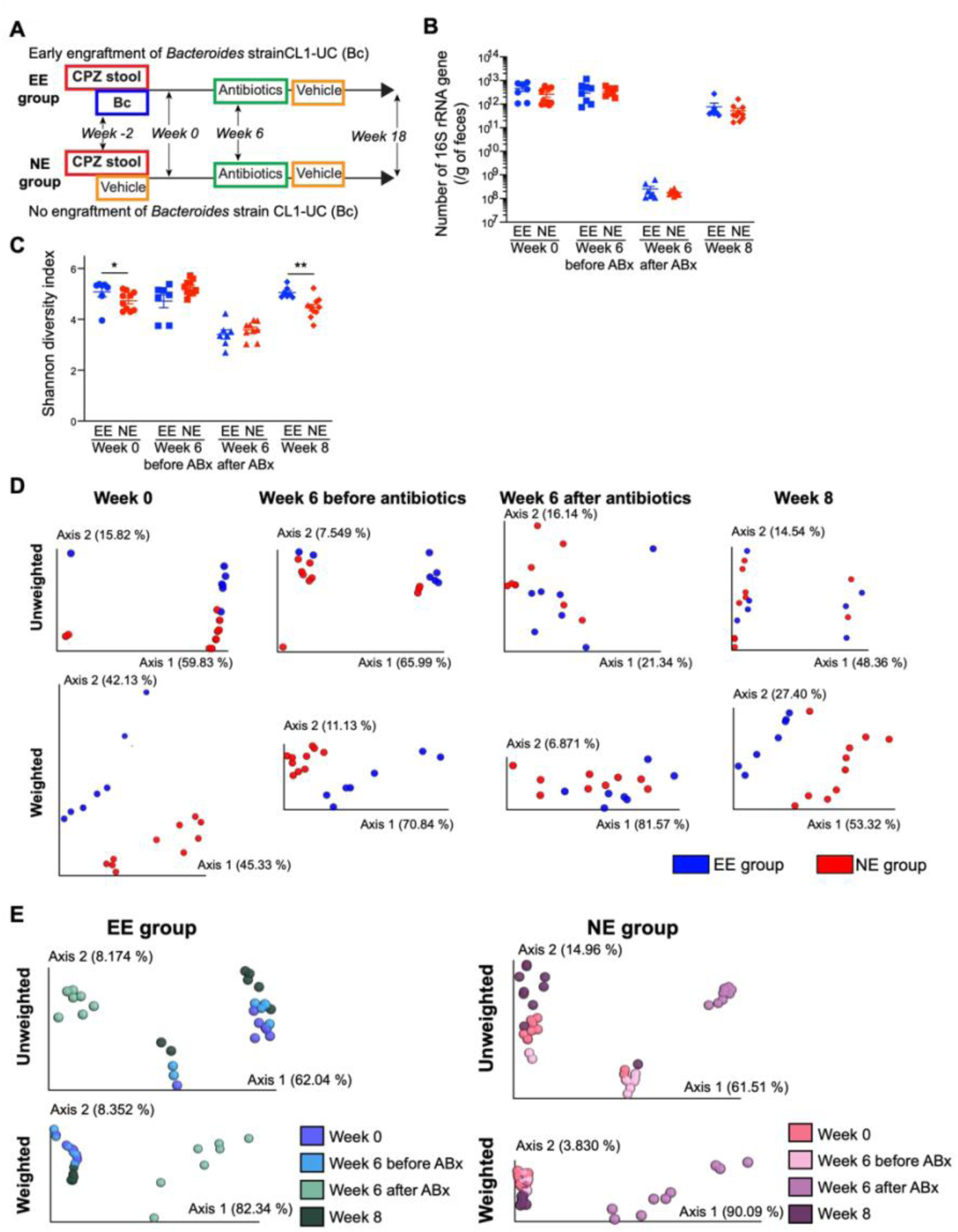
Early life engraftment of a single *Bacteroides* strain impacts bacterial membership of the gut microbiome. (A) Study design using the IL-10 knock-out mouse model. CPZ-induced dysbiosis was transplanted into germ-free pups at 3 weeks of age by fecal gavage. Following FMT, one cohort of mice were gavaged with the live *Bacteroides* strain CL1-UC (Bc) or vehicle control representing the early engraftment of Bc (EE) and no engraftment of Bc (NE) groups, respectively. (EE group: *n* = 7 and NE group: *n* = 10). Two weeks later (week 0; 5 weeks of age), animals started to be tracked. A cocktail of broad-spectrum antibiotics (neomycin, vancomycin, and CPZ) were administered at week 6 to perturb the gut microbiome and eradicate Bc in later stages of life, followed by observation to week 18. Details of the gavage and treatment schedules are presented in Figure S4. (B) The number of 16S rRNA gene per gram of feces of EE and NE animals over time (week 0, week 6 before antibiotics, week 6 after antibiotics, and week 8). (C) Shannon diversity index of EE and NE animals over time. (D) PCoA plots of unweighted and weighted UniFrac distances of 16S rRNA gene amplicon sequences between EE vs. NE groups at each time point (week 0, week 6 before antibiotics, week 6 after antibiotics, and week 8). The bacterial compositions were different between EE and NE groups at all time points. (E) PCoA plots of unweighted and weighted UniFrac distances showing over-time shifts of bacterial compositions in EE group (left panel) and NE group (right panel). The bacterial structure at week 8 recovered to a state before antibiotics treatment. \**p* < 0.05 and \*\**p* < 0.01. Data represent mean ± SEM for (B) and (C). See also Figure S4.

**Figure 5.**
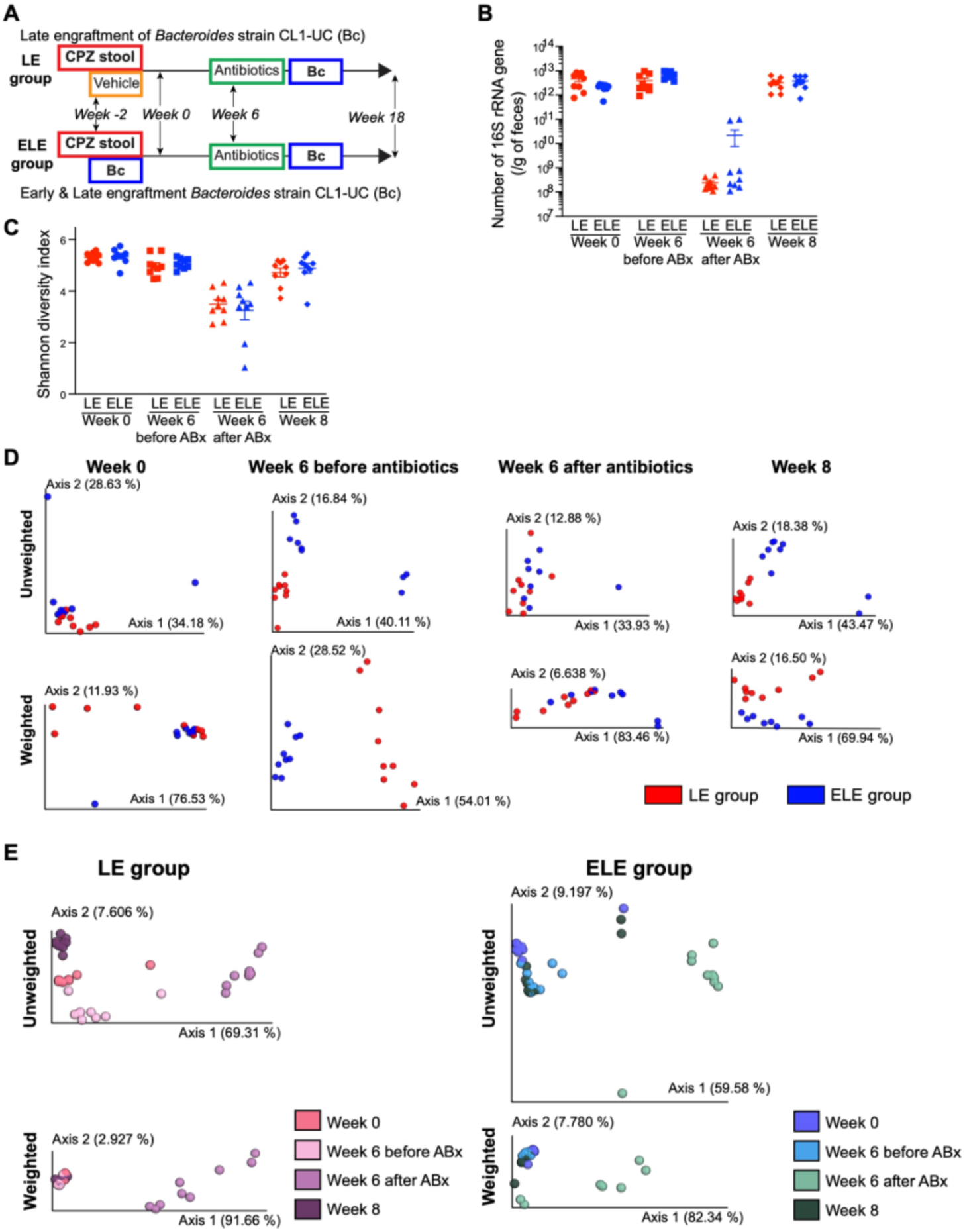
Later life engraftment of the *Bacteroides* strain does not impact bacterial membership of the gut microbiome. (A) Study design to determine the effects of late engraftment (LE) versus early and late engraftment (ELE) of Bc. The protocol is nearly identical to NE and EE groups shown in Figure 4, except that Bc is engrafted later in life in both LE and ELE groups (LE group: *n* = 9 and ELE group: *n* = 9). Late engraftment was performed at week 6 (11 weeks of age) after the administration of broad spectrum antibiotics cocktail meant to break microbiome stability and allow engraftment of Bc. Details of the gavage and treatment schedules are presented in Figure S5. (B) The number of 16S rRNA gene per gram of feces of LC and ELC animals over time (week 0, week 6 before antibiotics, week 6 after antibiotics, and week 8). (C) Shannon diversity index of LE and ELE animals over time. (D) PCoA plots of unweighted and weighted UniFrac distances of 16S rRNA gene amplicon sequences between LE vs. ELE groups at each time point (week 0, week 6 before antibiotics, week 6 after antibiotics, and week 8). The bacterial compositions were different between LE and ELE groups at all time points. (E) PCoA plots of unweighted and weighted UniFrac distances showing the gut microbiota composition shifts over time in LE group (left panel) and ELE group (right panel). Despite the antibiotic treatment and late engraftment of Bc, the bacterial structure at week 8 recovered toward that before antibiotics treatment. \**p* < 0.05 and \*\**p* < 0.01. Data represent mean ± SEM for (B) and (C). See also Figure S5.

We next investigated the impact of Bc engraftment on CPZ-associated gut microbiota over time at week 0 (2 weeks after initial gavage; T1), week 6 (before Abx treatment; T2), week 6 after Abx treatment (prior to the second gavage; T3), and week 8 (2 weeks after the second gavage; T4). No differences in 16S rRNA gene copy number were observed between EE and NE groups at all time points. After Abx exposure, the number of 16S rRNA gene dramatically decreased but recovered 2 weeks later (Figure 4B). Following Abx treatment, Shannon diversity index decreased but recovered 2 weeks after cessation, along with 16S rRNA gene copy number (Figure 4C). Unweighted and weighted UniFrac distances showed differences in community membership between EE and NE at all time points (T1: unweighted *p* = 0.05 and weighted *p* = 0.001, T2: unweighted *p* < 0.05 and weighted *p* = 0.001, T3: unweighted *p* < 0.05 and weighted *p* = 0.121, T4: unweighted *p* < 0.05 and weighted *p* = 0.001) (Figure 4D). The shift in relative abundance of the *Bacteroides* genus over time arose from engraftment and eradication of Bc in EE as compared to NE (Figure S4D). NE mice did not contain *Bacteroides*, while the presence of *Bacteroides* due to Bc engraftment was detected in EE mice at week 0 and week 6 before Abx. Bacterial composition at the phylum level in EE and NE groups at each time point is shown in Figure S4E. The *Bacteroides* genus corresponded to 59.7% and 53.2% of the Bacteroides phylum at week 0 and week 6 in the EE group before antibiotics. The phyla Bacteroidetes as well as the genus *Bacteroides* was eradicated in EE mice after Abx particularly at week 8, whereas mice in the NE group contained bacteria belonging to the phylum Bacteroidetes but not to the genus *Bacteroides*. Importantly, 2 weeks after cessation of Abx, community composition was indistinguishable from that observed in pre-treatment samples in both the EE and NE groups, respectively (Figure 4E). These findings indicate that engraftment with a single bacterial strain, such as Bc, early in life elicits a lasting impact on gut microbiome development and host immune tolerance to commensal microbes.

We next sought to test whether engraftment late in life of key microbes like Bc into CPZ-associated gut dysbiosis outside the window of immunological development can shift gut microbiota, restore immune homeostasis, and lower risk of spontaneous colitis in IL-10 KO mice. To this end a modification to the NE and EE group protocol (Figure 4A) was added where late life engraftment of Bc was performed at week 6 following Abx perturbation of the gut microbiome when mice were 11weeks of age (see Figure 5A). An Abx “shock” protocol is absolutely required to allow engraftment of a single organism, such as Bc into a stable microbial community where pre-existing priority conditions prevent non-resident community members from joining (Shen et al., 2015; Zmora et al., 2018). A cocktail of vancomycin, neomycin, and CPZ was administered in this study (details shown in Figure S5A). In order to compare the effects of early vs. late Bc engraftment, one group of mice received Bc gavage both early and late in life (at 3 weeks of age and 11 weeks of age) [designated early and late engraftment of Bc or ELE group] and the other where only late engraftment was performed at 11 weeks of age [late engraftment of Bc (LE) group]. This was done to determine impact of late Bc engraftment on additional gut microbial community members and immune homeostasis where mice were either pre-exposed (immune-tolerized) to Bc in early life or were Bc-naïve at the time of late Bc engraftment. Early life engraftment of Bc was confirmed in the ELE group 1 week after the first gavage at weaning (3 weeks of age) (Figure S5B) and again 1 week after Abx and Bc gavage at 11 weeks of age (Figure S5C). Bc also successfully engrafted in the LE group 1 week after cessation of Abx treatment (Figure S5C). Similar to our observations in the EE and NE groups, the estimated bacterial load and Shannon diversity were reduced by Abx but recovered after 2 weeks in the LE and ELE groups (Figures 5B and 5C). Bacterial composition in the LE and ELE groups were examined via 16S rRNA gene amplicon sequencing over time at week 0 (2 weeks after initial gavage; T1), week 6 (before Abx treatment; T2), week 6 after Abx treatment (prior to the second gavage; T3), and week 8 (2 weeks after the second gavage; T4). Unweighted and weighted UniFrac distances were significantly different between the LE and ELE groups at all time points (T1: unweighted *p* < 0.01 and weighted *p* = 0.133, T2: unweighted *p* = 0.001 and weighted *p* = 0.001, T3: unweighted *p* < 0.01 and weighted *p* < 0.01, T4: unweighted *p* = 0.001 and weighted *p* < 0.05) (Figure 5D). Shifts in the relative abundances the genus *Bacteroides* members over time arose from engraftment and eradication of Bc in LE and ELE mice (Figure S5D). The presence of the *Bacteroides* genus due to Bc engraftment was detected in LC mice at week 8. In ELE mice, early and late engraftment of Bc was confirmed by presence of the *Bacteroides* genus at weeks 0 and 8. Bacterial composition at the phylum level in the LE and ELE groups at each time point is shown in Figure S5E. LE mice contained bacteria belonging to the Bacteroides phylum but not the *Bacteroides* genus before late engraftment, while the *Bacteroides* genus corresponded to 99.7% of Bacteroides phylum at week 8. In ELE mice, the genus *Bacteroides* corresponded to 97.8%, 49.0%, 27.5%, and 58.1% of the phylum Bacteroides at week 0, week 6 before Abx, week 6 after Abx, and week 8, respectively. Despite Bc engraftment later life following Abx treatment, bacterial composition at T4 was similar to T1 and T2 in the LE and ELE groups, respectively (Figure 5E).

### Colonization with a single Bacteroides strain early in life mitigates colitis development

We next examined the impact of early life Bc engraftment on development of spontaneous colitis in the IL-10 KO cohorts described above. EE mice exhibited increased weight gain compared to their NE group counterparts (Figure 6A). None of the EE group animals developed clinical symptoms of colitis, while 2 out of 10 NE group mice reached the euthanasia criteria due to rectal prolapse and severe loss of body weight during the observation period (Figure 6B). At week 6 (11 weeks of age), but before antibiotic treatment, fecal LCN-2 levels were significantly elevated in NE mice relative to the EE group (*p* < 0.05) (Figure 6C). Fecal LCN-2 level remained elevated in NE mice as compared to EE mice at week 18 (*p* < 0.05), the observational endpoint (23 weeks of age) (Figure 6C). Similarly, LE mice exhibited elevated levels of fecal LCN-2 compared to ELE mice at week 6 before Abx exposure (*p* < 0.001) as well as at week 18 (*p* < 0.05) (Figure 6D), although no animals in the LE and ELE groups reached the euthanasia criteria during the observation period. Overall, early life colonization with Bc can prevent spontaneous colitis despite its eradication later life via Abx, while colonization with Bc later in life does not elicit a beneficial effect.

**Figure 6.**
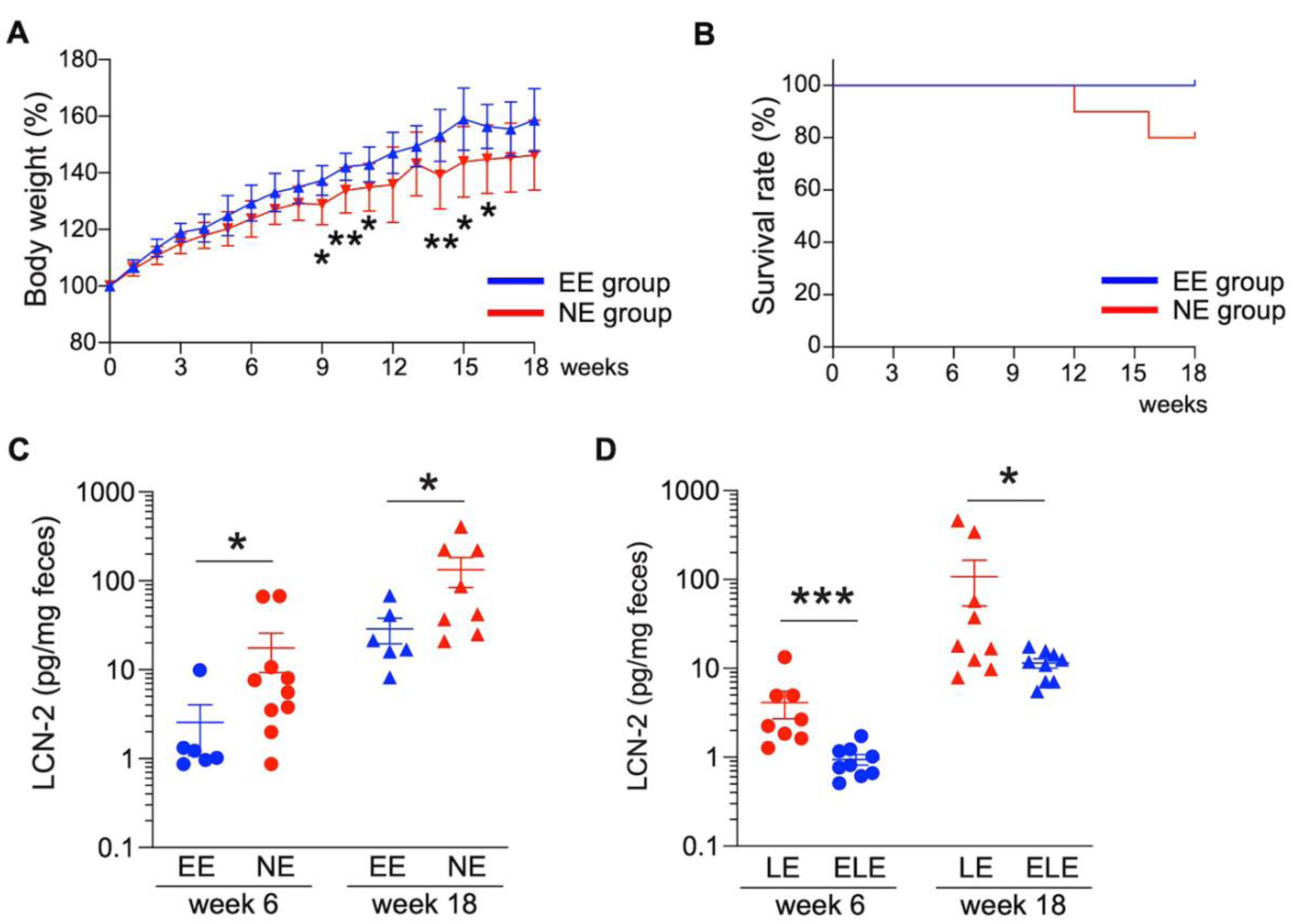
Engraftment of a single *Bacteroides* strain in early life prevents colitis development. (A) Percent weight change (expressed as % of starting weight at week 0) of the tracked animals beginning at week 0 (5 weeks of age). (B) Survival rate of animals in EE and NE groups during the 18-week observation period. Two out of ten animals in NE group reached the euthanasia criteria. (C) Fecal lipocalin-2 (LCN-2) levels before antibiotic treatment (week 6) and at the end point of the observation period (week 18) in EE and NE animals. (D) Fecal lipocalin-2 (LCN-2) levels before antibiotics treatment (week 6) and at the end point of the observation period (week 18) in LE and ELE animals. \**p* < 0.05, \*\**p* < 0.01, and \*\*\**p* < 0.001. Data are represented as mean ± SEM for (A), (C), (D).

## Discussion

Several findings emerged from our studies that could have broad implications to our current understanding of human IBD etiopathogenesis and in rethinking clinical best practices. Our findings support that perturbations in the early life gut microbiome can have negative and persistent consequences to immune development that lead to increased risk for disease in genetically susceptible hosts. Moreover, our past and current findings strongly suggest that these events arise from a “loss of function”, i.e. extinction of critical CPZ-sensitive commensal microbes in the maternal gut microbiota that are not vertically transmitted to offspring, rather than a “gain of function”, i.e. emergence of pathobionts that activate the immune system of genetically susceptible hosts to trigger the onset of disease. In support of this notion, when IL10-deficient GF dams and sires are conventionalized with CPZ-induced donor gut microbiota, reproduce, and continue to be housed in the same gnotobiotic isolator as their offspring, neither dams, sires, nor offspring develop spontaneous colitis. In contrast, offspring from CPZ-treated dams housed in SPF conditions appear to re-acquire certain CPZ-sensitive commensal gut microbiota from the environment later in life and develop spontaneous colitis (Miyoshi et al., 2017). These microbes, in an immunologically-naïve/insufficiently-tolerized host are thought to induce an immune response that increases risk for development of spontaneous colitis. We identified and cultivated a unique indigenous *Bacteroides* strain (*Bacteroides* strain CL1-UC, Bc) using sequence-based and bioinformatics approaches from fecal samples obtained from offspring of CPZ-treated dams that developed spontaneous colitis later in life (Figure 3). Bc successfully engrafted in recipients at 3 weeks of age that had vertically-transmitted CPZ-induced perturbed microbiota. The early engraftment of Bc significantly impacts the ensuing gut bacterial assemblage and reduces risk for development of spontaneous colitis, but these effects were observed only if the intervention was made early in life (Figures 4, 6, and S4). Engraftment of Bc into CPZ-altered gut microbiota of IL-10 KO offspring later in life required a perturbation with a brief course of antibiotics. We believe this is because of the priority effects of established gut microbiota, which has already undergone a selection process to establish community stability and resilience. Under these conditions, “foreign” microbes, even if commensal under other circumstances, would be excluded from becoming part of the community (Shen et al., 2015; Zmora et al., 2018). A short course of antibiotics was therefore needed to “reset” community dynamics to allow Bc to engraft (Figures 5 and S5). The finding that early, but not late engraftment, confers an IL-10-independent immune tolerance further supports the paradigm for the necessity of immunological conditioning or imprinting that only occurs in a developmental window. Future studies are needed to determine whether early exposure to Bc elicits a generalized immune tolerance to either novel bacterial or environmental antigen exposure later in life that prevents disease onset. Together, by testing hypotheses emerged from multi-omics observations through cultivation and colonization experiments, our findings support the notion that perturbations in early life immune and gut microbiome development can significantly affect subsequent states of health and disease and that best practices should include measures to identify host-microbiome imbalances and means to correct them at this critical life stage. These might include a focus on early life risk stratification and intervention, particularly for complex immune disorders that have been increasing in incidence and prevalence with alarming frequency over the past century. Significant perturbations of the gut microbiome are caused by factors like antibiotic exposure, diet, e.g. breast versus formula feeding, birth modality, e.g., vaginal vs. Caesarian section, and environment that contribute to loss of critical microbes and mediators essential to this process (Sonnenburg et al., 2016). The resulting negative consequences can potentially set the stage for long term disease risk, especially in hosts genetically prone to complex immune disorders such as IBD.

Current modern medicine practices are rooted in identifying events temporally related to disease onset and then to intervene after-the-fact. At this point, many additionally amplifying and self-sustaining processes are set into play, particularly in genetically susceptible individuals, that result in chronic and often medically refractory disease. Our study further suggests that interventions to correct microbial imbalances must be done during an early life stage, as they may become less effective once the immune conditioning period is over. The success of any microbiome-based intervention or biotherapeutic hinges on defining the early life “healthy” gut microbiome. This remains a challenge with few solutions, as the gut microbiota of individuals varies considerably at the taxonomical level. However, by identifying microbial populations and their derived mediators that provide specific beneficial functions in a defined context such as immune development in early life could be a plausible solution. We have provided proof-of-concept that introduction of a single indigenous member of the gut microbiota (Bc) at the right time and in the right context can lower disease risk. On a practical level, it is unlikely that this particular bacteria or other murine-derived microbes will play the same role, exhibit identical biological properties, or possess the ability to engraft into human subjects because the assembly rules for acquisition and colonization of gut microbes by each host species differs (Rawls et al., 2006; Turnbaugh et al., 2009). However, similar strategies to ours could be modified to identify and develop human-specific microbes and their mediators that have biotherapeutic potential. In doing so, the knowledge gained may aid in developing new metrics and biomarkers for defining and assessing the healthy gut microbiome and in stratifying individuals who would receive the greatest benefit from early microbiome-based interventions. With this, a suite of next generation microbiome-based therapeutics are likely to emerge that are effective for both disease prevention as well as treatment of complex diseases.

## Materials and Methods

### Resource availability

Further information and requests for resources and reagents should be directed to and will be fulfilled by the Lead Contact, Eugene B. Chang (echang at medicine dot bsd dot uchicago dot edu).

### Materials Availability

Frozen stock of Bacteroides strain CL1-UC isolated and cultivated in the present study is stored in the facility of the Lead Contact.

### Data Availability

We have made publicly available raw sequencing data for all metagenomes (see Table S1 for MG-RAST IDs). In addition, as anvi’o profiles we have made available (1) individual co-assemblies and the final set of metagenome-assembled genomes (doi:10.6084/m9.figshare.11955060) which offer reproducible access to detection and coverage statistics of each genome across metagenomes and (2) pangenomes (doi:10.6084/m9.figshare.11968674 for Figure 3B and doi:10.6084/m9.figshare.11968692 for Figure 3C) which offer reproducible access to functions and sequences of each gene cluster and their distribution across genomes. The raw sequencing data for 16S rRNA gene amplicon sequences reported in this study is also publicly available (MG-RAST mgp82768).

### Animals

All mice used for these studies were on a C57Bl/6J genetic background. Specific pathogen free (SPF) mice were bred in the University of Chicago animal vivarium under Helicobacter hepaticus-free conditions (Institutional Animal Care and Use Committee protocol 71084). Germ-free IL-10 KO mice and RAG1KO mice were bred in the University of Chicago Gnotobiotic Research Animal Facility (GRAF).

### Peripartum antibiotic treatment

The peripartum CPZ treatment protocol was identical to our previous study (Miyoshi et al., 2017). To normalize the gut microbiota among breeding pairs in no treatment (NT) and CPZ groups, prior to onset of the study, a weekly bedding transfer protocol (Miyoshi et al., 2018) was performed for sires and dams from 3 weeks of age (weaning) until 8 weeks of age (when breeders were paired). Pups from these breeders were sacrificed and tissues were harvested at 7 weeks of age (Figure 1A) (Institutional Animal Care and Use Committee protocol 72348).

### T cell transfer study

Five germ-free RAG1 KO breeding pairs were conventionalized with H. hepaticus-free microbiota from C57Bl/6J wild-type mice animals bred at the University of Chicago. The weekly bedding transfer protocol was performed for their progeny until breeding was initiated. Pups from these normalized breeding pairs were used as RAG1 KO recipients at 8-10 weeks of age. Sex-matched NT and CPZ IL-10 KO mice at 7 weeks of age were used as donors. CD4^+^ T cells were isolated from SPLNs of 5 donors from each group using a mouse CD4^+^ T cell isolation kit following manufacturer’s instructions (STEMCELL Technologies, Vancouver, BC). CD4^+^ T cells from NT and CPZ donors were injected intraperitoneally into RAG1 KO recipient mice, respectively (1.0 × 10^6^ cells in 500 μL of sterile PBS). Control RAG1 KO recipients were injected with 500 μL sterile PBS. Disease progression monitoring and colitis assessment were continued until 8 weeks post-injection (Ostanin et al., 2009). The euthanasia criteria included greater than 20% body weight loss, rectal prolapse, frank blood in the stool, and decreased food and water consumption (Institutional Animal Care and Use Committee protocol 72348).

### Fecal microbiota transplantation and engraftment of Bacteroides strain CL1-UC in gnotobiotic conditions

GF IL-10 KO male mice were used to examine the impact of Bacteroides strain CL1-UC (Bc) engraftment on colitis development. Fecal samples collected from 7-week-old NT and CPZ male pups from our previous study (Miyoshi et al., 2017) were used as donors for gavage of 100 μL fecal solution (100 mg feces per 1 ml sterile PBS) into GF IL-10 KO male recipients at 3 weeks of age (day 0 of early engraftment). For early engraftment of Bc at 3 weeks of age, recipients were gavaged 100 μL (because of the smaller body size) of bacterial solution (5 × 10^8^ CFU/ml in PBS) on days 1 and 2 (Figure S4A). Animals without early engraftment were gavaged 100 uL with sterile PBS alone. Bacterial colonization was confirmed, and tracking of pups began 2 weeks post-conventionalization (week 0). At week 6 (11 weeks of age), pups were treated with a broad spectrum antibiotic cocktail [vancomycin (0.5 mg/ml), neomycin (1.0 mg/ml), and CPZ (0.5 mg/ml) in the drinking water] for 72 hours (days 0-2 of late engraftment) and allowed a 24-hour recovery period (day 3). For late engraftment, animals were gavaged 200 μL of bacterial solution (5 × 10^8^ CFU/ml) on days 4 and 5 (Figure S5A). Animals without late engraftment were gavaged 200 μL of sterile PBS alone. We assessed early engraftment (EE) vs. no engraftment (NE) groups and late engraftment late engraftment (LE) vs. early and late engraftment (ELE) groups. Each of the 4 groups was maintained in an individual semi-rigid containment isolator to prevent the influence of external environmental microbes for the duration of the study from weaning until 23 weeks of age. The euthanasia criteria for spontaneous colitis were identical to those previously described (Miyoshi et al., 2017) including rectal prolapse, more than 15% body weight loss compared to maximum body weight, or signs of pain/distress, including poor grooming, decreased activity, and hunched posture (Institutional Animal Care and Use Committee protocol 72101).

### Fecal lipocalin-2 measurement

Frozen fecal samples were diluted in lysis buffer [100mg feces per 1 ml of PBS containing protease inhibitors (Roche, Indianapolis, IN)]. Fecal lysates were analyzed using a mouse lipocalin-2/NGAL ELISA kit following the manufacturer’s instructions (R&D Systems, Minneapolis, MN).

### Histological analysis

samples were fixed in 4% formaldehyde and embedded in paraffin followed by H&E staining. The histological score for colitis in the CD4^+^ T cell transfer model was assessed as previously reported (Erben et al., 2014).

### Fecal DNA extraction and 16S rRNA gene amplicon sequencing

Fecal samples were harvested and rapidly frozen at −80°C. Fecal DNA extraction and sequencing were performed using previously reported protocols (Miyoshi et al., 2017). Briefly, the V4 region of the 16S rRNA gene was amplified following the EMP protocol (http://www.earthmicrobiome.org/protocols-and-standards/16s/) and sequences were obtained by Illumina MiSeq at the Argonne Sequencing Center (Lemont, IL). The raw sequencing data were analyzed by QIIME2 2018.8 (Bolyen et al., 2018) with DADA2 (Callahan et al., 2016). Samples with less than 4,000 sequences were excluded from the analyses. The taxonomy was analyzed using the taxonomy classifier trained on the Greengenes 13_8 99% OTUs that is provided by QIIME2 development team (https://docs.qiime2.org/2019.4/data-resources/). We used qPCR to calibrate the number of 16S rRNA gene per gram of feces.

### Shotgun metagenomics and genome-resolved metagenomics

Metagenomic shotgun sequencing for fecal DNA samples was performed on Illumina HiSeq at the Marine Biological Laboratory (Woods Hole, MA) which yielded paired-end reads of 2×150 nucleotides. Noisy sequences were removed using ‘iu-filter-quality-minoche’ (Eren et al., 2013) with default parameters, a program that implements noise filtering parameters (Minoche et al., 2011). The remaining reads were co-assembled using MEGAHIT (Li et al., 2015) v1.0.3 with a minimum scaffold length of 2.5 kbp, and scaffold header names were simplified using anvi’o (Eren et al., 2015) v6.1 (available from https://merenlab.org/software/anvio). Scaffolds were subsequently binned as previously described (Delmont et al., 2018; Eren et al., 2015). Briefly, (1) an anvi’o contigs database was generated from assembled scaffolds using ‘anvi-gen-contigs-database’ program where during which Prodigal (Hyatt et al., 2010) v2.6.3 with default parameters identified open reading frames, and HMMER (Eddy, 2011) v3.1b2 identified genes matching to bacterial single-copy core genes (Campbell et al., 2013); (2) the program ‘anvi-import-taxonomy-for-genes’ imported gene-level taxonomic annotations predicted by Centrifuge (Kim et al., 2016) against NCBI’s nt database into anvi’o; (3) short reads were mapped to the scaffolds for each metagenome using Bowtie 2 (Langmead and Salzberg, 2012) v2.0.5 and the recruited reads were stored as BAM files using SAMtools (Li et al., 2009); (4) ‘anvi-profile’ was used to process each BAM file for each sample to estimate the coverage and detection statistics of each scaffold, and combined mapping statistics were stored in an anvi’o merged profile database using ‘anvi-merge’. Scaffolds were then clustered with the automatic binning algorithm CONCOCT (Alneberg et al., 2014) using the program ‘anvi-cluster-contigs’ with the parameter ‘--clusters 25’ to constrain the number of bins to 25 prior to the final manual refinement step. Finally, we used ‘anvi-refine’ to manually refine each CONCOCT bin using the anvi’o interactive interface. Bins with <10% of redundancy, and with either >2Mbp in length or >70% of estimated completion were defined as metagenome-assembled genomes (MAGs), manually curated using the same interactive interface. The program ‘anvi-summarize’ reported final FASTA files for MAGs and CheckM (Parks et al., 2015) to infer their taxonomy using 43 single-copy gene markers. The anvi’o merged profile database that gives reproducible access to MAGs and their distribution across samples is accessible at doi:10.6084/m9.figshare.11955060.

### Pangenome analysis

To compute a pangenome of MAGs and isolates we used the anvi’o pangenomic workflow as previously described (Delmont and Eren, 2018). Briefly, this approach consists of two major steps: (1) generating an anvi’o genome database using the program ‘anvi-gen-genomes-storage’ (v.6.1) to store DNA and amino acid sequences, as well as functional annotations of each gene in each genome, (2) and computing the pangenome using the program ‘anvi-pan-genome’. During the second step ‘gene clusters’ are identified by first determining sequence similarities between genes with a reciprocal search using the NCBI’s BLASTP program (Altschul et al., 1990) and later resolving the network of associations using the MCL algorithm (van Dongen and Abreu-Goodger, 2012) with default parameters. We also used the program ‘anvi-compute-genome-similarity’ to calculate average nucleotide identity between genomes, which relied on PyANI (Pritchard et al., 2016) and import the heatmap into the final pangenome. Results were then visualized in the anvi’o interactive interface using the program ‘anvi-display-pan’ and ‘anvi-summarize’ gave access to gene cluster distribution patterns across MAGs and isolates and gene sequences in gene clusters. The URL http://merenlab.org/p describes the details of the anvi’o pangenomics workflow. The anvi’o pan database that gives reproducible access to gene clusters across genomes is accessible at doi:10.6084/m9.figshare.11968674 for Figure 3B and doi:10.6084/m9.figshare.11968692 for Figure 3C

### Whole genome sequencing for bacterial DNA and assembling genomes

Bacterial DNA was extracted from a single colony using a microbial DNA extraction kit (QIAGEN, Germantown, MD). Whole genome sequencing for bacterial DNA was performed on Illumina MiSeq at Marine Biological Laboratory (Woods Hole, MA). The raw sequencing data were assembled with PATRIC (Wattam et al., 2014).

### Isolation and cultivation of bacterial strain corresponding to target metagenomic-assembled genome

Fecal samples were homogenized in sterile PBS under anaerobic conditions (200 mg/ml). The solutions were diluted to 10^−5^ and streaked on anaerobic Brucella agar plates with 5% sheep blood, hemin and vitamin K_1_ (BD, Sparks, MD) with 0.1mg/ml of kanamycin. Individual colonies were screened for taxonomy based on 16S rRNA gene sequencing for each colony at the University of Chicago Genomics Facility using universal primers as previously described. PCR was performed for candidate colonies using primers designed for each target MAG. These selected and characterized isolates were cultured in Brucella broth (BD, Sparks, MD) with hemin (5 μg/mL) and Vitamin K (0.0001%) and 20% glycerol (5 × 10^9^ CFU/mL) stocks were prepared for storage at −80°C.

### Statistical analysis

Mann-Whitney U test was performed with GraphPad Prism (GraphPad Software, CA, USA) to compare 16s rRNA gene copy number, Shannon diversity index, T cell populations, body weights, histological scores, fecal LCN-2 levels, and cytokine levels between NT and CPZ groups. For 16S rRNA gene amplicon sequencing analysis, PERMANOVA tests were computed with QIIME2 (Bolyen et al., 2018) to assess bacterial composition differences between treatment groups or for unweighted and weighted UniFrac distances data. Statistical significance was achieved at p < 0.05. MAG detection rates between CPZ-colitis group vs. CPZ-no-colitis group at 11 weeks of age were compared using student’s t-test and the p-values were adjusted for multiple-test using the Benjamini-Hochberg method (Benjamini and Hochberg, 1995). The criterion for significance was set at a false discovery rate (FDR) < 0.05.

### Flow cytometry

Spleens (SPLNs) and mesenteric lymph nodes (MLNs) were harvested for assessing CD4^+^ T cell populations and cytokine production of CD4^+^ T cells. FcR blocking was performed for all samples with anti-mouse CD16/CD32 antibody (BD Biosciences, CA, San Jose). Cells were stained using the LIVE/DEAD Fixable Aqua Dead Cell Stain Kit (Thermo Fisher Scientific, Waltham, MA) to assess viability. For surface stains, anti-mouse TCRβ (eBioscience, San Diego, CA) and anti-mouse CD4 antibodies (Biolegend, San Diego, CA) were used. For intranuclear stains, the Foxp3/Transcription Factor Staining Buffer Set (eBioscience) was used followed by staining with anti-mouse Foxp3 (eBioscience), anti-mouse T-bet (eBioscience) and anti-RORγt antibodies (eBioscience). For cytokines, anti-mouse IFNγ antibody (Biolegend) and anti-IL-17A antibody (Thermo Fisher Scientific) were used. The Rat IgG2a Kappa Isotype, Mouse IgG1 Kappa Isotype, and Rat IgG1 Kappa Isotype controls (eBioscience) were used. Samples were analyzed using a FACSCanto (BD Biosciences) and FlowJo v10 (FLOWJO, OR, Ashland).

### Dendritic cell stimulation assay with fecal lysates

DCs were isolated from SPLNs of mice at 7 weeks of age using a mouse CD11c positive selection kit II (STEMCELL Technologies, Vancouver, BC). Fecal lysates used for stimulation were prepared by resuspending fecal samples (100 mg feces in 1ml of PBS) followed by bead-beating, centrifugation, and recovery of the supernatant. DCs (5 × 10^5^ cells/well, 1ml of the medium/well) were incubated at 37°C for 24 hours with 0.1 nM retinoic acid (Sigma-Aldrich, St. Louis, MO) and 2 ng/ml recombinant mouse TGF-β (R&D Systems, Minneapolis, MN).

### CD4^+^ T cell cytokine production assay

For stimulation with PMA/ionomycin, cells were obtained from SPLNs and MLNs. Red blood cells lysis was performed on SPLNs. Cells (2.5 × 10^6^ cells/well, 1 ml of medium/well) were stimulated with PMA (50 ng/ml) and ionomycin (500 ng/ml) with Golgistop (1.5ul/ml) (BD Biosciences) for 3 hours at 37°C with 5% CO_2_. Cells were then harvested for flow cytometry to examine IFNγ and IL-17A production. For stimulation with anti-CD3/anti-CD28 antibodies, cell culture plates were pre-coated with anti-mouse CD3 antibody (Biolegend) (1 μg/ml) and anti-mouse CD28 antibody (BD Biosciences) (2 μg/ml). CD4^+^ T cells were harvested from SPLNs and MLNs using a CD4^+^ T cell isolation kit (STEMCELL technologies). Cells (2 × 10^5^ cells/well, 200 μl of the medium/well) were incubated at 37°C with 5% CO_2_ for 72 hours. The supernatants were collected to analyze cytokine production via ELISA.

### ELISA

The concentrations of IL-12p40, IFNγ, and IL-17A were measured in frozen supernatant samples obtained from cell culture outlined above. Commercially available ELISA kits for these mouse cytokines were used following the manufacturer’s instructions (Thermo Fisher Scientific).

### Bacterial culture and bacterial lysates

Bacteroides strain CL1-UC (Bc), B. thetaiotaomicron (Bt), Lactobacillus murinus (Lm), and Lachnospiraceae bacterium (Lb) were cultured to obtain bacterial lysates for immunological experiments. Bc and Lm were isolated from murine feces in our laboratory. Bt isolated from mice is a kind gift from Prof. Alexander Chervonsky (University of Chicago). Lb was purchased from ATCC (ATCC BAA-2281). In anaerobic conditions, Bc was cultured in Brucella broth with hemin and vitamin K, Bt was cultured in BHI broth, Lm was cultured in MRS broth, and Lb was cultured in modified chopped meat media. Bacteria were grown to mid-log or stationary phase (O.D. = 0.6-0.9), followed by centrifugation and bacterial pellets were washed with sterile PBS. The bacterial pellets were resuspended in PBS and bead-beaten. After centrifuging, supernatants were collected. The protein concentrations of these lysates were determined by BCA assay and 400 μg/ml stock solutions were prepared for immunological experiments.

### Dendritic cell stimulation assay and co-culture of dendritic cell/naïve CD4^+^ T cell with bacterial lysates

were isolated from SPLN and incubated as described above. For DC stimulation with bacterial lysates for 24 hours, the final protein concentration added to each well was 20 μg/ml in the incubation media. DCs and naïve CD4^+^ T cells were harvested from SPLNs of mice at 7 weeks of age using a CD11c positive selection kit II (STEMCELL Technologies) and a mouse naïve CD4^+^ T cell Isolation kit (STEMCELL Technologies). The cell culture plates were pre-coated with anti-mouse CD3 antibody (Biolegend) (1 μg/ml). DCs (4 × 10^4^ cells/well) and naïve CD4^+^ T cells (6 × 10^4^ cells/well) were plated together with 0.1 nM retinoic acid (Sigma-Aldrich) and 2 ng/ml recombinant mouse TGF-β (R&D Systems). Cells were incubated with bacterial lysates at 37°C for 72 hours. The final protein concentration of the lysates was 20 μg/ml in the incubation media. After 72-hour incubation, supernatants and cells were harvested to examine cytokine production and T cell differentiation.

## Author Contributions

J.M., S.M., V.L., A.M.E., and E.B.C. conceived the study, designed experiments, and prepared the manuscript. J.M., S.M., C.C., A.S., K.Y., and Y.S. performed experiments and analyzed data. E.K., M.Y., and A.M.E. provided analysis tools. J.M., T.O.D., S.T.M.L., D.A.A., and A.M.E. analyzed microbial data sets. V.L., S.C., M.S., D.A.A., and E.B.C. supervised the manuscript. E.B.C. and V. L. oversaw the entire project.

## Acknowledgments

The present research was supported by the NIDDK Digestive Disease Core Research Center (NIH P30 DK42086); NIDDK grants R37 DK47722 (to E.B.C.) and K01 DK111785 (V.L.); the University of Chicago GI Research Foundation. We acknowledge generous support from the David and Ellen Horing Research Fund. We thank the Human Tissue Resource Center for histological processing (the University Chicago), the Gnotobiotic Research Animal Facility staff for GF animal husbandry (the University of Chicago) as well as the Flow Cytometry Core Facility for analysis software (Kyorin University Graduate School of Medicine). We also thank Mark W. Musch for sample acquisition/analysis.

**Tables and Supplementary Figures**

**Figure S1. Maternal peripartum antibiotic exposure induces persistent gut microbiota perturbations and a chronic imbalance of CD4**^**+**^ **T cell subsets in female offspring**.

(A) The number of 16S rRNA gene per gram of feces in female offspring from non-treated (NT) and CPZ-exposed dams at 3 and 7 weeks of age (*n* = 5 in each group). (B) Shannon diversity index in female offspring at 3 and 7 weeks of age. (C) PCoA plots of unweighted and weighted UniFrac distances of 16S rRNA gene amplicon sequences in NT and CPZ female pups at 3 and 7 weeks of age, respectively. (D) Bar charts represent relative abundances of phyla in fecal samples from NT and CPZ pups at 3 and 7 weeks of age. (E) Representative images of flow cytometry gating strategies for analyzing T cell populations with representative isotype controls. (F) Flow cytometric analyses of live CD4^+^ T cells expressing Foxp3 (Treg), T-bet (Th1) or RORγt (Th17) in spleens (SPLNs) and mesenteric lymph nodes (MLNs) of NT versus CPZ females at 7 weeks of age (*n* = 4-5 per group). Data represent the percentage of live TCRβ^+^CD4^+^ cells. (G) IL-12p40 production of dendritic cells obtained from 7-week-old female NT (blue) and CPZ (red) pups with stimulation by fecal samples (*n* = 5 per group). \*\**p* < 0.01. Data represent mean ± SEM for (A), (B), (F), and (G).

**Figure S2. Maternal peripartum antibiotic exposure promotes a proinflammatory state of CD4**^**+**^ **T cells in female offspring**.

(A) Flow cytometric analyses of live CD4^+^ T cells from offspring expressing IFNγ and IL-17A with stimulation via PMA and ionomycin. Cells from spleen (SPLN) and mesenteric lymph nodes (MLNs) were obtained from NT and CPZ female pups at 7 weeks of age (*n* = 4-5 per group). Data represent the percentage of live TCRβ^+^CD4^+^ cells. (B) Representative images of flow cytometry gating strategies for analyzing T cell populations with representative isotype controls. (C) IFNγ and IL-17A production of CD4^+^ T cells obtained from SPLN and MLN of 7-week-old NT and CPZ female pups with co-stimulation via anti-CD3 and anti-CD28 antibodies. (D) The number of 16S rRNA gene per gram of feces of female RAG1 KO recipients before CD4^+^ T cell transfer (NT recipients: *n* = 13, CPZ recipients: *n* = 14, Control group: *n* = 12). (E) Shannon diversity index of female recipients before CD4^+^ T cell transfer. (F) PCoA plots of unweighted and weighted UniFrac distances of 16S rRNA gene amplicon sequences in female recipients before CD4^+^ T cell transfer. (G) Percent weight change (expressed as % of starting weight) of female recipients after CD4^+^ T cell transfer. (H) Survival rate of female recipients during the 8-week observation period. (I) Fecal lipocalin-2 (LCN-2) levels before CD4^+^ T cell transfer (week 0) and 2 weeks post-transfer (week 2) in female recipients. (J) Histological assessment of colons from surviving female recipients at week 8 (NT recipients: *n* = 12 and CPZ recipients: *n* = 13). Representative H&E histological sections of colon are presented for NT recipients, CPZ recipients, and the control group. \**p* < 0.05, \*\**p* < 0.01, and \*\*\**p* < 0.001. Data represent mean ± SEM for (A), (C)-(E), (G), (I), and (J).

**Figure S3. Isolation of *Bacteroides* strain CL1-UC cultivar and assessment of its immunological impact**

(A) Validation of primers targeted towards identified metagenomic-assembled genomes (MAGs 1-10) in fecal samples. Four primer sets were designed for each MAG 1-10. The fecal sample positive for these MAGs were used to validate these primers. NC: negative control, containing 16S rRNA primers without fecal DNA. PC: positive control, the fecal sample plus 16S rRNA gene primers. (B) Comprehensive genome analysis via PATRIC performed phylogenetic analysis showing our strain, *Bacteroides* strain CL1-UC, was closely related to Bacteroides sp. CAG:754 and reference organism Bacteroides finegoldii DSM 17565, but unique enough to be considered a novel strain. (C) IL-12p40 production of dendritic cells (DCs). DCs were isolated from spleen of IL-10 KO mice at 7 weeks of age and were stimulated using bacterial lysates from the following strains; *Bacteroides* strain CL1-UC(Bc), *B. thetaiotaomicron* (Bt), *Lactobacillus murinus* (Lm), and Lachnospiraceae bacterium (ATCC BAA-2281) (Lb), or BLNK: blank control with sterile PBS (*n* = 5 per group). (D) IL-12p40 and IFNγ production and Th1 differentiation in the co-culture assay of DCs and naïve CD4^+^ T cells. Flow cytometry data represent the percentage of live TCRβ^+^CD4^+^T-bet^+^ cells out of live TCRβ^+^CD4^+^ cells (*n* = 4-5 per group). \**p* < 0.05, \*\**p* < 0.01, and \*\*\**p* < 0.001. Data represent mean ± SEM for (C) and (D).

**Figure S4. Design of the CPZ-induced gut microbiota conventionalization protocol with timed delivery of *Bacteroides* strain CL1-UC early in life with corresponding confirmation of engraftment via PCR and 16S rRNA gene amplicon sequencing**.

(A) Study design of fecal conventionalization and bacterial delivery via gavage. Four to 5 mice were housed per cage for both the early engraftment (EE) group with *Bacteroides* strain CL1-UC (Bc) and no colonization (NE) group (*n =* 9 to 10 mice per group). EE and NE animals were maintained in separate semi-rigid isolators throughout the study, respectively. See also Figure S5. (B) Fecal samples obtained from 2 mice per cage (4 mice in total from each group) were used to verify Bc engraftment via PCR one week after early gavage. (C) PCR for Bc detection was performed using fecal samples from these same animals one week after the second sterile PBS gavage. Negative controls did not contain fecal DNA template. (D) Relative abundance of genus *Bacteroides* in EE and NE groups over time (week 0, week 6 before antibiotics, week 6 after antibiotics, and week 8) based on 16S rRNA gene amplicon sequences. NE group did not contain *Bacteroides*, while the presence of *Bacteroides* due to Bc engraftment was detected in EE group at week 0 and week 6 before antibiotics. (E) Bar charts represent relative abundances of phyla in EC and NC groups over time (week 0, week 6 before antibiotics, week 6 after antibiotics, and week 8) based on 16S rRNA gene amplicon sequences. The bacterial compositions at week 8 recovered toward that observed before antibiotics treatment.

**Figure S5. Design of CPZ-induced gut microbiota conventionalization protocol with timed delivery of *Bacteroides* strain CL1-UC early and late in life with corresponding confirmation of engraftment via PCR and 16S rRNA gene amplicon sequencing**.

(A) Study design of fecal conventionalization and bacterial delivery via gavage for late engraftment (LE) and early and late engraftment (ELE). Four to 5 mice were housed per cage for both the LE group and ELE group (*n =* 9 to 10 mice per group). LE and ELE animals were maintained in separate semi-rigid isolators throughout the study, respectively. See also Figure S4. (B) Fecal samples obtained from 2 mice per cage (4 mice in total from each group) were used to verify Bc engraftment via PCR one week after early gavage. (C) PCR for Bc detection was performed using fecal samples from these same animals one week after the last gavage. Negative controls did not contain fecal DNA template. (D) The relative abundance of genus *Bacteroides* in LE and ELE groups over time (week 0, week 6 before antibiotics, week 6 after antibiotics, and week 8) based on 16S rRNA gene amplicon sequences. The presence of *Bacteroides* genus due to Bc engraftment was detected in LC group at week 8. In ELE group, early and late engraftment of Bc was confirmed by the presence of *Bacteroides* genus at weeks 0 and 8. (E) Bar charts represent phylum relative abundances in LE and ELE groups over time (week 0, week 6 before antibiotics, week 6 after antibiotics, and week 8) based on 16S rRNA gene amplicon sequences. The bacterial compositions at week 8 recovered to a state that was observed before antibiotic treatment.

