## Supplementary figures and images for "Early-life microbial intervention reduces colitis risk promoted by antibiotic-induced gut dysbiosis"

### Supplementary Figure S1

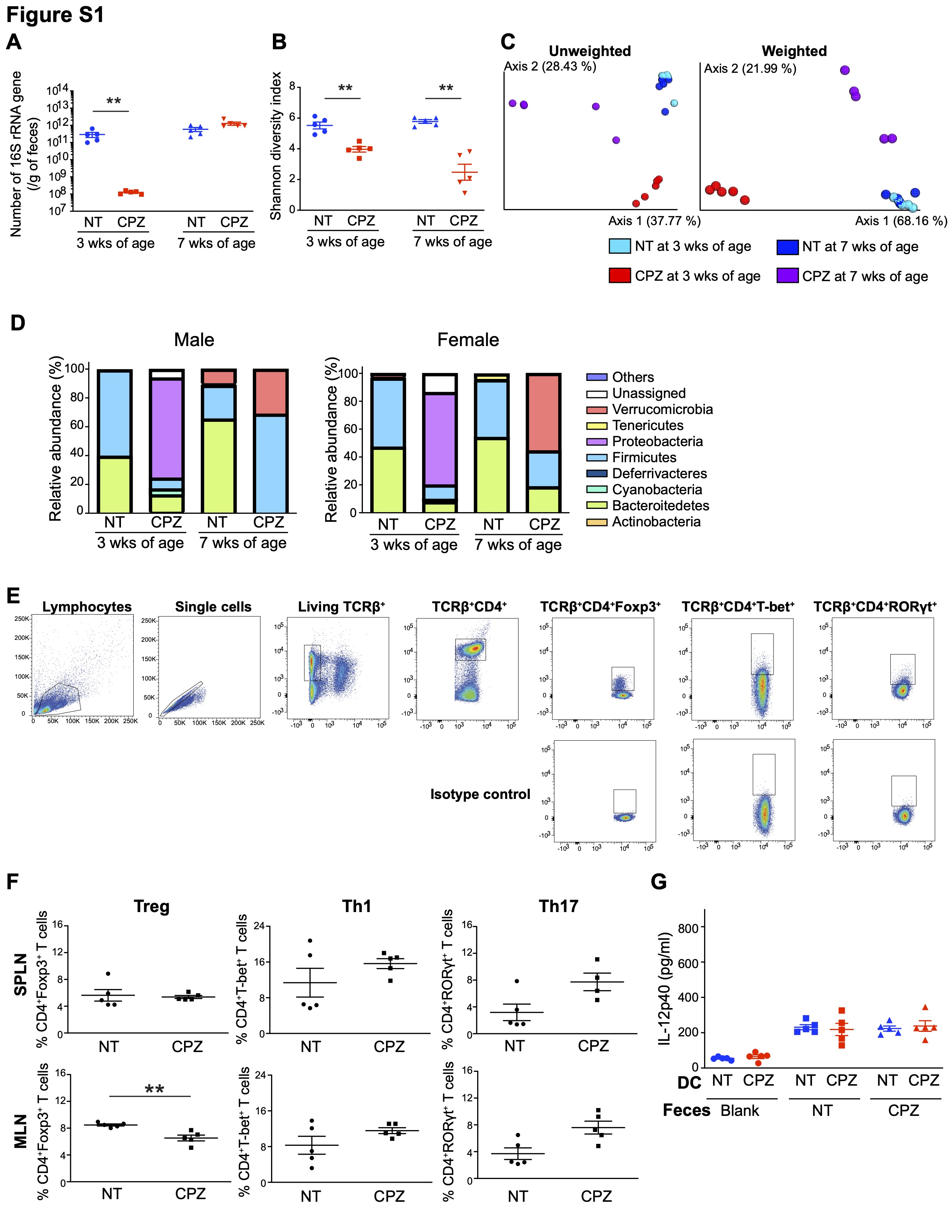

### Supplementary Figure S2

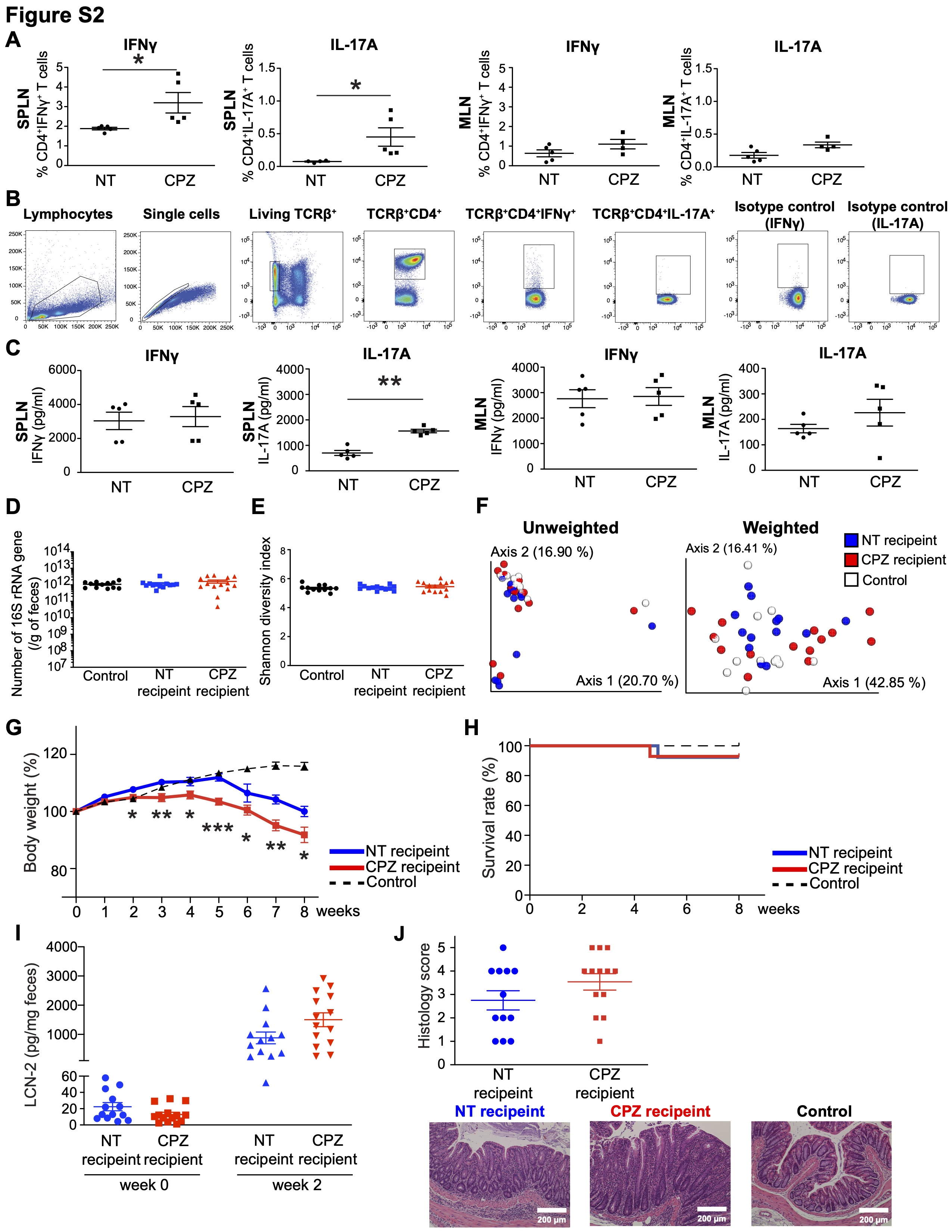

### Supplementary Figure S3

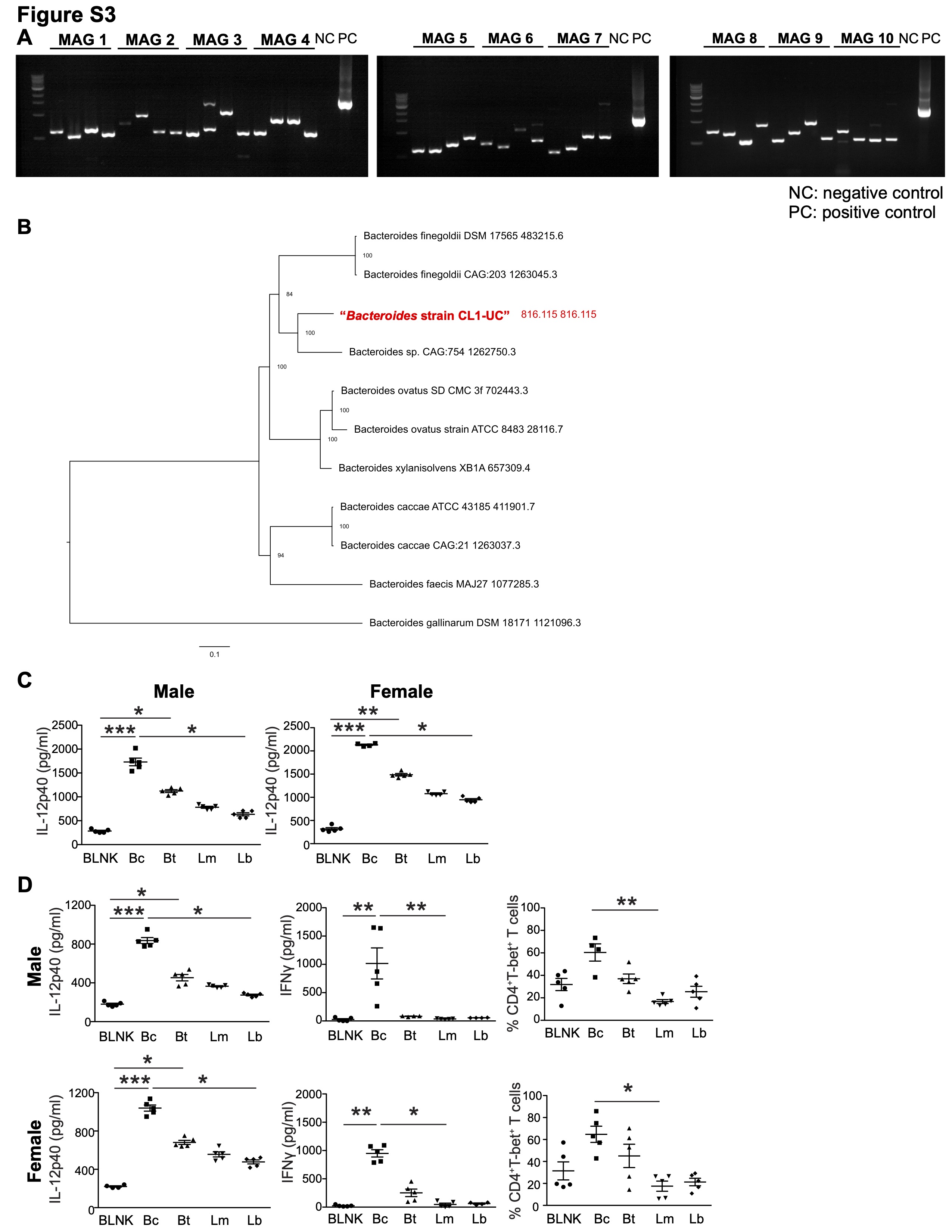

### Supplementary Figure S4

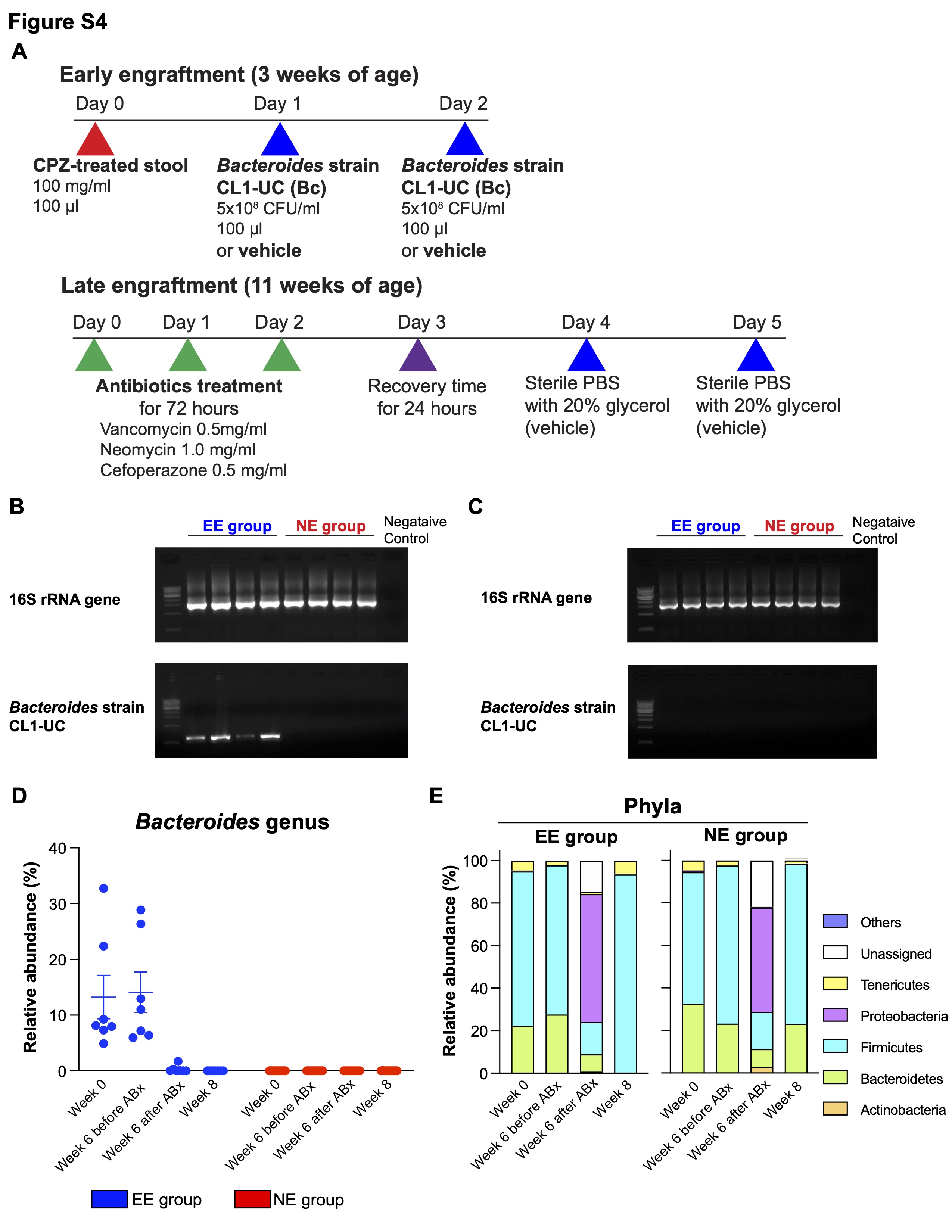

### Supplementary Figure S5

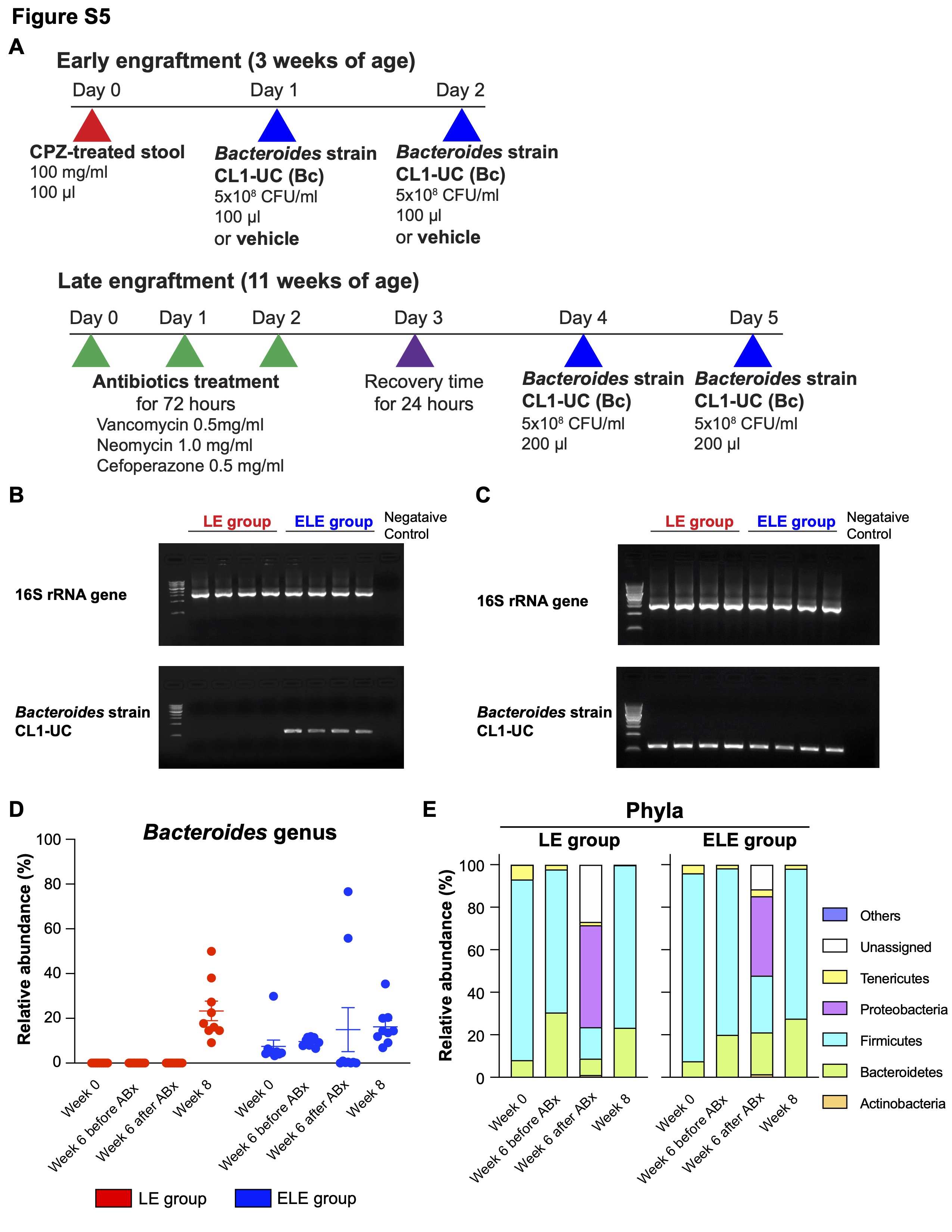
