## Supplementary Table S2 for "Early-life microbial intervention reduces colitis risk promoted by antibiotic-induced gut dysbiosis"

Table S2. Metagenome-assembled genomes (MAGs) 1-10 and reference MAGs for anvi’o pangenome analysis

|  | **Comparison of MAG detection rate**  **between CPZ-colitis group (Col) vs. CPZ-no-colitis group (Non)**  **at 11 weeks of age** | |
| --- | --- | --- |
| **MAG 1** | Col > Non; | Female: *p* < 0.05 & Male: *p* < 0.05 |
| **MAG 2** | Col > Non; | Female: *p* < 0.05 & Male: *p* < 0.05 |
| **MAG 3** | Col > Non; | Female: *p* < 0.05 & Male: *p* < 0.05 |
| **MAG 4** | Col > Non; | Female: *p* < 0.05 & Male: *p* < 0.05 |
| **MAG 5** | Col > Non; | Female: *p* < 0.05 & Male: *p* < 0.05 |
| **MAG 6** | Col > Non; | Female: *p* < 0.05 & Male: *p* < 0.10 |
| **MAG 7** | Col > Non; | Female: *p* < 0.05 & Male: *p* < 0.05 |
| **MAG 8** | Col > Non; | Female: *p* < 0.05 & Male: *p* < 0.05 |
| **MAG 9** | Col > Non; | Female: *p* < 0.05 & Male: *p* < 0.05 |
| **MAG 10** | Col > Non; | Female: *p* < 0.05 & Male: *p* < 0.05 |
| **Reference 1** | Col > Non; | Female: *p* < 0.05 |
| **Reference 2** | Col < Non; | Male: *p* < 0.05 |
| **Reference 3** | Col < Non; | Male: *p* < 0.05 |
| **Reference 4** | Col > Non; | Male: *p* < 0.05 |
| **Reference 5** | Col > Non; | Female: *p* < 0.05 & Male: *p* < 0.05 |
| **Reference 6** | Col < Non; | Male: *p* < 0.05 |
| **Reference 7** | Col < Non; | Female: *p* < 0.05 |
| **Reference 8** | Col > Non; | Male: *p* < 0.05 |
| **Reference 9** | Col < Non; | Male: *p* < 0.05 |
| **Reference 10** | Col < Non; | Female: *p* < 0.05 |
