## Supplementary Table S3 for "Early-life microbial intervention reduces colitis risk promoted by antibiotic-induced gut dysbiosis"

Table S3. Primers for metagenome-assembled genomes (MAGs) 1-10

|  |  | **Forward** | **Reverse** |
| --- | --- | --- | --- |
| **MAG1** | 1 | TGTTGCCTTGATCGTCTCTG | TACCAATACTCCCCCATCCA |
|  | 2 | AACCCTCACCGTACATCAGC | TCCGCAAAAGGCTGTAGAGT |
|  | 3 | CCGTAAATGGCTGCGATGTG | GAATGATGCAGCGGGAGAGA |
|  | 4 | CAACAGTTCGAGTTGGCTGA | GCCGTTTGGAACATCGTACT |
| **MAG2** | 1 | GGAACATACGTGAGGCGTTT | GCATCTCCTGCTGGCTAATC |
|  | 2 | CCCGGTGTAAGCGGTACTAA | TCCGACTCATATCCGTAGCC |
|  | 3 | TGGAATAACCGTCCTATGCT | TATTCCTTCAGGCAGGCAAC |
|  | 4 | AGGTTACCTCTGCGGGATCT | CGTGGAAAACCTCAACCAGT |
| **MAG3** | 1 | CATCCGCTCTGAACAAGACA | GAGTTTGTCGGCAGGGAATA |
|  | 2 | GCTGCCTGTCATCTGCTGTA | AAGCATCAAAGACCGGATTG |
|  | 3 | CGGAAGCGTTCTTTCTGTTC | TATACCGACGCCGAGATACC |
|  | 4 | ACGGTTATAAGGCGGAGGTC | GCACAGTGAAAACACCGAAA |
| **MAG4** | 1 | TCGATATCTGCGTGAACAGC | CGAAAACTGCCAAACAGACA |
|  | 2 | CGCCCTTGGATTATCTGAAA | AAGAGGTTTGTCGGGAAGGT |
|  | 3 | GACTATTTCCGGGTGGAGGT | GTAGACCGACCGCATATCGT |
|  | 4 | GGTTATTGTCGTGCATGTCG | GTCGCGATAAACAGTCAGCA |
| **MAG5** | 1 | TTGTCAAGGCCCTCGATTAC | GAGGCACAACGCTGTACAAA |
|  | 2 | AGCAGCGCATAGGCTATCAT | ACGATGAGGGTCGTTACAGG |
|  | 3 | ATGTCGACCGTGGATTCTTC | GCTCGTTGAGCGTTATGTCA |
|  | 4 | TCGAGGTTGTGAGCTTCCTT | AGGGCACAAAGCTTGAAGAA |
| **MAG6** | 1 | TTGTACGGGTTTCCTTCGAC | ATGTCGCAGCGTCCTATCTT |
|  | 2 | TTATCAACGGTCGGCCTTAC | CAGGCCCAAGGTACTCACAT |
|  | 3 | GAGCGGAGTCTGTCGTTTTC | TCCGGCTTCGTTGTTTTATC |
|  | 4 | CAAAACATTCGGCATACACG | TAGGGGCTGTGGGTTGTAAG |
| **MAG7** | 1 | CGCGGTACTGCTGACAGATA | CATACAGCTGGTAGGCAGCA |
|  | 2 | AATGGTTTGCCAATGCTTTC | TGATTCGGTGGTATGGGATT |
|  | 3 | GTCCTTTTTCGCGCTCTATG | CCGTTGTTCGGTCTGAATTT |
|  | 4 | TACAGATACCCCGGAACTGC | GATACAAACGGCAACGGTCT |
| **MAG8** | 1 | GACGATGACGGGATCACTCT | CCATTGACGTAACCGAAGGT |
|  | 2 | AGTCACAGCGGCTTCAAACT | TGTCCTGCCAGACAACAGAG |
|  | 3 | TTGAAACGCATCTCCTTTCC | ATGGCAGTAAGCACGACTTG |
|  | 4 | TCACGATCAAAGACGTCAGC | GCTGTTCATGCTTTCGTCAA |
| **MAG9** | 1 | AACGCTCTCCTCGTGAAAAA | AACTCTGTGGCGGTGGATAC |
|  | 2 | GGCTCTCGATGGGATTGATA | GCTACACCAGTCCCGACATT |
|  | 3 | GCGAAATCCACATAGCGTTT | TAGCGATGCCAACAAGACAG |
|  | 4 | TTCATTGCCGAATACGACCT | CCGGTACTATGACCGACGAT |
| **MAG10** | 1 | CTAAACAACCGGCATCCACT | ACGTCTGATGGGAAAGTTGG |
|  | 2 | AATTTCACCACCGTCCAGAG | GGCCTTGATGTCCAGAAAAA |
|  | 3 | GCAACTGTCGACCTGACGTA | CCAAGTCTGTTTCAGGCACA |
|  | 4 | ATTTACTGACGGGCCAAGTG | GGACGGCAAGCGTATAACAT |
