## Supplementary Table S4 for "Early-life microbial intervention reduces colitis risk promoted by antibiotic-induced gut dysbiosis"

Table S4. Percentages of Average Nucleotide Identity (ANI) between MAG 1, Isolates 1-18, and reference bacterial genomes

|  | **Ref. 1** | **Ref. 2** | **Ref. 3** | **Ref. 4** | **MAG 1** | **Isolate 1** | **Isolate 2** | **Isolate 3** | **Isolate 4** | **Isolate 5** | **Isolate 6** | **Isolate 7** | **Isolate 8** | **Isolate 9** | **Isolate 10** | **Isolate 11** | **Isolate 12** | **Isolate 13** | **Isolate 14** | **Isolate 15** | **Isolate 16** | **Isolate 17** | **Isolate 18** |
| --- | --- | --- | --- | --- | --- | --- | --- | --- | --- | --- | --- | --- | --- | --- | --- | --- | --- | --- | --- | --- | --- | --- | --- |
| **Ref. 1** | 100.00 | 83.61 | 81.31 | 81.39 | 83.82 | 83.84 | 83.83 | 83.84 | 83.83 | 83.83 | 83.84 | 83.83 | 83.83 | 83.83 | 83.83 | 83.84 | 83.84 | 83.83 | 83.84 | 83.84 | 83.83 | 83.83 | 83.84 |
| **Ref. 2** | 83.52 | 100.00 | 81.94 | 81.55 | 89.17 | 90.08 | 90.06 | 90.06 | 90.08 | 90.09 | 90.08 | 90.07 | 90.08 | 90.07 | 90.07 | 90.08 | 90.07 | 90.08 | 90.06 | 90.08 | 90.07 | 90.08 | 90.07 |
| **Ref. 3** | 81.06 | 81.69 | 100.00 | 98.69 | 81.19 | 81.29 | 81.29 | 81.29 | 81.29 | 81.29 | 81.29 | 81.29 | 81.29 | 81.29 | 81.29 | 81.29 | 81.29 | 81.29 | 81.29 | 81.29 | 81.29 | 81.29 | 81.29 |
| **Ref. 4** | 81.15 | 81.31 | 98.56 | 100.00 | 81.20 | 81.26 | 81.26 | 81.26 | 81.26 | 81.26 | 81.26 | 81.26 | 81.26 | 81.26 | 81.26 | 81.26 | 81.26 | 81.26 | 81.26 | 81.26 | 81.26 | 81.26 | 81.26 |
| **MAG 1** | 83.88 | 89.48 | 81.39 | 81.47 | 100.00 | 100.00 | 100.00 | 100.00 | 100.00 | 100.00 | 100.00 | 99.99 | 100.00 | 100.00 | 100.00 | 100.00 | 100.00 | 100.00 | 100.00 | 100.00 | 100.00 | 100.00 | 100.00 |
| **Isolate 1** | 83.77 | 90.15 | 81.37 | 81.36 | 99.85 | 100.00 | 99.99 | 99.99 | 99.99 | 99.99 | 99.99 | 99.99 | 99.99 | 99.99 | 99.99 | 99.99 | 99.99 | 99.99 | 99.99 | 99.99 | 99.99 | 100.00 | 99.99 |
| **Isolate 2** | 83.75 | 90.16 | 81.37 | 81.36 | 99.83 | 99.99 | 100.00 | 99.99 | 99.99 | 99.99 | 99.99 | 99.99 | 99.99 | 99.98 | 99.99 | 99.99 | 99.99 | 99.99 | 99.99 | 99.99 | 99.99 | 99.99 | 99.99 |
| **Isolate 3** | 83.76 | 90.15 | 81.35 | 81.34 | 99.83 | 99.99 | 99.99 | 100.00 | 99.99 | 99.99 | 99.99 | 99.99 | 99.99 | 99.99 | 99.99 | 99.99 | 99.99 | 99.99 | 99.99 | 99.99 | 99.99 | 99.99 | 99.99 |
| **Isolate 4** | 83.75 | 90.10 | 81.35 | 81.35 | 99.84 | 99.98 | 99.98 | 99.98 | 100.00 | 99.98 | 99.98 | 99.98 | 99.98 | 99.98 | 99.98 | 99.98 | 99.98 | 99.98 | 99.98 | 99.98 | 99.98 | 99.98 | 99.98 |
| **Isolate 5** | 83.75 | 90.14 | 81.33 | 81.32 | 99.83 | 99.98 | 99.99 | 99.98 | 99.99 | 100.00 | 99.99 | 99.99 | 99.98 | 99.98 | 99.98 | 99.99 | 99.99 | 99.99 | 99.98 | 99.99 | 99.99 | 99.99 | 99.99 |
| **Isolate 6** | 83.79 | 90.17 | 81.40 | 81.39 | 99.85 | 99.99 | 99.99 | 99.99 | 99.99 | 99.99 | 100.00 | 99.99 | 99.99 | 99.99 | 99.99 | 99.99 | 99.99 | 99.99 | 99.99 | 99.99 | 99.99 | 99.99 | 99.99 |
| **Isolate 7** | 83.76 | 90.17 | 81.39 | 81.38 | 99.84 | 99.99 | 99.99 | 99.99 | 99.99 | 99.99 | 99.99 | 100.00 | 99.99 | 99.98 | 99.99 | 99.99 | 99.99 | 99.99 | 99.99 | 99.99 | 99.99 | 99.99 | 99.99 |
| **Isolate 8** | 83.77 | 90.14 | 81.35 | 81.34 | 99.83 | 99.99 | 99.99 | 99.99 | 99.99 | 99.99 | 99.99 | 99.98 | 100.00 | 99.99 | 99.99 | 99.99 | 99.99 | 99.99 | 99.99 | 99.98 | 99.99 | 99.99 | 99.99 |
| **Isolate 9** | 83.77 | 90.15 | 81.37 | 81.34 | 99.85 | 99.99 | 99.99 | 99.99 | 99.99 | 99.99 | 99.99 | 99.99 | 99.99 | 100.00 | 99.99 | 99.99 | 99.99 | 99.99 | 99.99 | 99.99 | 99.99 | 99.99 | 99.99 |
| **Isolate 10** | 83.77 | 90.17 | 81.40 | 81.38 | 99.85 | 99.99 | 99.99 | 99.99 | 99.99 | 99.99 | 99.99 | 99.99 | 99.99 | 99.99 | 100.00 | 99.99 | 99.99 | 99.99 | 99.99 | 99.99 | 99.99 | 99.99 | 99.99 |
| **Isolate 11** | 83.76 | 90.12 | 81.37 | 81.36 | 99.84 | 99.99 | 99.99 | 99.99 | 99.99 | 99.99 | 99.99 | 99.98 | 99.99 | 99.99 | 99.99 | 100.00 | 99.99 | 99.99 | 99.99 | 99.99 | 99.99 | 99.99 | 99.99 |
| **Isolate 12** | 83.78 | 90.15 | 81.38 | 81.36 | 99.84 | 99.99 | 99.99 | 99.99 | 99.99 | 99.99 | 99.99 | 99.99 | 99.99 | 99.98 | 99.99 | 99.99 | 100.00 | 99.99 | 99.99 | 99.99 | 99.99 | 99.99 | 99.99 |
| **Isolate 13** | 83.75 | 90.12 | 81.35 | 81.34 | 99.84 | 99.99 | 99.99 | 99.99 | 99.99 | 99.99 | 99.99 | 99.99 | 99.99 | 99.99 | 99.99 | 99.99 | 99.99 | 100.00 | 99.99 | 99.99 | 99.99 | 99.99 | 99.99 |
| **Isolate 14** | 83.77 | 90.13 | 81.35 | 81.33 | 99.84 | 99.99 | 99.99 | 99.99 | 99.99 | 99.99 | 99.99 | 99.99 | 99.99 | 99.99 | 99.99 | 99.99 | 99.99 | 99.99 | 100.00 | 99.99 | 99.99 | 99.99 | 99.99 |
| **Isolate 15** | 83.74 | 90.12 | 81.37 | 81.36 | 99.84 | 99.99 | 99.99 | 99.99 | 99.99 | 99.99 | 99.99 | 99.99 | 99.99 | 99.99 | 99.99 | 99.99 | 99.99 | 99.99 | 99.99 | 100.00 | 99.99 | 99.99 | 99.99 |
| **Isolate 16** | 83.76 | 90.13 | 81.39 | 81.38 | 99.84 | 99.99 | 99.99 | 99.99 | 99.99 | 99.99 | 99.99 | 99.99 | 99.99 | 99.99 | 99.99 | 99.99 | 99.99 | 99.99 | 99.99 | 99.99 | 100.00 | 99.99 | 99.99 |
| **Isolate 17** | 83.75 | 90.13 | 81.35 | 81.34 | 99.84 | 99.99 | 99.99 | 99.99 | 99.99 | 99.99 | 99.99 | 99.99 | 99.99 | 99.99 | 99.99 | 99.99 | 99.99 | 99.99 | 99.99 | 99.99 | 99.99 | 100.00 | 99.99 |
| **Isolate 18** | 83.77 | 90.10 | 81.34 | 81.32 | 99.84 | 99.99 | 99.99 | 99.99 | 99.99 | 99.99 | 99.99 | 99.99 | 99.99 | 99.99 | 99.99 | 99.99 | 99.99 | 99.99 | 99.99 | 99.99 | 99.99 | 99.99 | 100.00 |

Ref. 1: *Bacteroides caccae* ATCC43185, Ref. 2: *B. caecimuris* I48, Ref. 3: *B. thetaiotaomicron* 7330, Ref. 4: *B. thetaiotaomicron* VPI5482
